# Airborne environmental DNA for terrestrial vertebrate community monitoring

**DOI:** 10.1101/2021.07.16.452634

**Authors:** Christina Lynggaard, Mads Frost Bertelsen, Casper V. Jensen, Matthew S. Johnson, Tobias Guldberg Frøslev, Morten Tange Olsen, Kristine Bohmann

## Abstract

Assessing and studying the distribution, ecology, diversity and movements of species is key in understanding environmental and anthropogenic effects on natural ecosystems. Although environmental DNA is rapidly becoming the tool of choice to assess biodiversity ^1–3^ there are few eDNA sample types that effectively capture terrestrial vertebrate diversity and those that do can be laborious to collect, require special permits and contain PCR inhibitory substances, which can lead to detection failure. Thus there is an urgent need for novel environmental DNA approaches for efficient and cost-effective large-scale routine monitoring of terrestrial vertebrate diversity. Here we show that DNA metabarcoding of airborne environmental DNA filtered from air can be used to detect a wide range of local vertebrate taxa. We filtered air at three localities in Copenhagen Zoo, detecting mammal, bird, amphibian and reptile species present in the zoo or its immediate surroundings. Our study demonstrates that airDNA has the capacity to complement and extend existing terrestrial vertebrate monitoring methods and could form the cornerstone of programs to assess and monitor terrestrial communities, for example in future global next generation biomonitoring frameworks ^4,5^.

## Main

Biodiversity monitoring at the community scale is a critical element of assessing and studying species distributions, ecology, diversity and movements ^e.g. 6,7^. Further, it informs conservation efforts, evaluates status and quotas on species subject to recreational or commercial harvest, detects the arrival of invasive species, and tracks progress in achieving biodiversity targets; crucial aims in light of the current climate and biodiversity crisis ^7–9^. This highlights the urgent need for efficient and cost-effective methods with which to document and monitor biological communities.

Over the last decade the analysis of environmental DNA, or eDNA, has emerged as a valuable tool for non-invasive, sensitive and cost-effective characterization of biodiversity and species communities that complements and extends existing methods ^1–3^. Typically eDNA is extracted from samples such as sediments, water, faeces or gut contents, and is a complex mixture of intra- and extracellular DNA derived from many sources and of different qualities ^1^. DNA metabarcoding coupled with high-throughput sequencing is generally used to sequence taxonomically informative markers ^10^. This has allowed compilation of species inventories, detection of common, rare, indicator and invasive species, and has provided information about plant-pollinator interactions and ecosystem services and dynamics ^e.g. 11–17^. Further, there is progress towards implementation of eDNA in routine biodiversity monitoring at both local and global scales ^4,5,18,19^.

Vertebrates are key species in most terrestrial ecosystems, but are experiencing extinctions and declines in population numbers and sizes due to increasing threats from human activities and environmental change ^20–23^ ; www.iucnredlist.org. Terrestrial vertebrate monitoring is, however, generally expensive, laborious and difficult with existing methods, and so far, terrestrial vertebrate monitoring with eDNA has been challenged by that only few of the currently applied eDNA sample types are capable of capturing community-scale terrestrial vertebrate diversity. Two eDNA sample types dominate such analyses: freshwater samples and invertebrate gut contents. In freshwater, terrestrial vertebrates can be detected through the DNA they leave when e.g. drinking or defecating ^24–28^, and DNA from vertebrates can be detected in the gut contents of parasitic, scavenging or coprophagous invertebrates ^13,29–32^.

However, invertebrate and freshwater samples can require permits and be laborious to collect, and may contain enzyme inhibitors such as heme compounds and humic acids which can hinder or introduce stochasticity in the metabarcoding PCR amplification of vertebrate DNA, leading to false negatives ^33–35^. Further, they represent relatively biased samples of vertebrate DNA due to potential invertebrate feeding preferences ^36^ and bias towards terrestrial vertebrates leaving DNA in freshwater ^28^. Hence, for eDNA-based monitoring of terrestrial vertebrates there is a gap between the operational difficulties and shortcomings of the currently established substrates and the urgent need for innovative, efficient and cost-effective methods for assessing vertebrate community composition.

We hypothesised that DNA captured from the air could solve these issues, potentially allowing for straightforward collection and characterisation of community scale distribution data from terrestrial vertebrates. Air is filled with particles, such as fungal spores, bacteria, vira, pollen, dust, sand, droplets and fibrous material, which can be airborne for days and transported over long distances in the atmosphere depending on humidity and particle size ^37–42^. These contain DNA and/or carry DNA attached to them, and recently DNA sequencing has been used to identify the taxonomic origins of airborne fungal spores, algae, pollen and microbiota collected on adhesive tape, in air filters and in dust traps ^43–49^. Further, two studies have indicated that micro-sized tissue fragments and debris from vertebrates can be airborne and detected through DNA-sequencing. One study demonstrated vertebrate detection from DNA filters in a small, confined room containing hundreds of individuals of the target species ^50^. Another study sequenced DNA from atmospheric dust samples in the Global Dust Belt over the Red Sea and detected eukaryotes, including small sequence quantities of human, cetacean and bird ^51^. However, the use of airDNA for studying and monitoring local vertebrate communities in a wider context is unexplored. Here, we demonstrate that a wide range of local terrestrial vertebrate taxa can be detected by sequencing of particles filtered from air, providing a new framework for airDNA assessment of terrestrial vertebrate communities.

### Terrestrial vertebrates leave detectable DNA in air

To investigate whether terrestrial vertebrates leave detectable DNA traces in air, we filtered air in Copenhagen Zoo, Denmark, which provided an ideal, controlled setting with a well-defined population of vertebrates exotic to the surrounding environment. Air was filtered for between 30 mins and 30 hrs using three different samplers; a water vacuum using line power in which air circulated through sterile water which was then filtered using a Sterivex filter, and two air particle samplers with class F8 fibrous filters for airborne particulate matter. One used a 24 V blower fan requiring line power and to pave the way for vertebrate monitoring in the wild, the other used a 5 V blower fan with a mobile phone power bank. Each sampling was carried out in duplicate: two consecutive replicates for the water vacuum, and two simultaneous replicates for each of the samples taken with the novel particle samplers, resulting in a total of 40 samples across the three sampling locations. DNA was extracted and high-throughput sequenced at two mtDNA metabarcoding markers: one targeting vertebrates in general and the other mammals specifically ^52,53^. In our data analysis we only retained Operational Taxonomic Units (OTU) that could be identified at species level, thereby providing a conservative inventory of vertebrate detections.

We first tested airDNA monitoring in a well-ventilated semi-confined space by collecting 12 airborne particulate matter samples in a stable in the southern section of the zoo holding two okapis (*Okapia johnstoni*) and two red forest duikers (*Cephalophus natalensis*) (Fig. 1a). Using this approach, we detected both the species present in the stable in all of the 12 samples. Further, we detected 13 birds and mammals that are kept in neighbouring outdoor enclosures in the southern section of the zoo, 1 zoo animal that was located in the northern section of the zoo, 2 animals kept in the zoo but that are also known to be pests, 2 wild or domestic non-zoo species known to occur in and around the zoo, and 2 fish species used as feed in the zoo (Fig. 2). Thus, overall, we detected 22 non-human vertebrate species (Fig. 2; Supplementary Table 1) with the number of species detected per sample ranging from 6 to 17 (mean = 11.33, SD= 3.17) (Supplementary Table 2).

**Figure 1.**
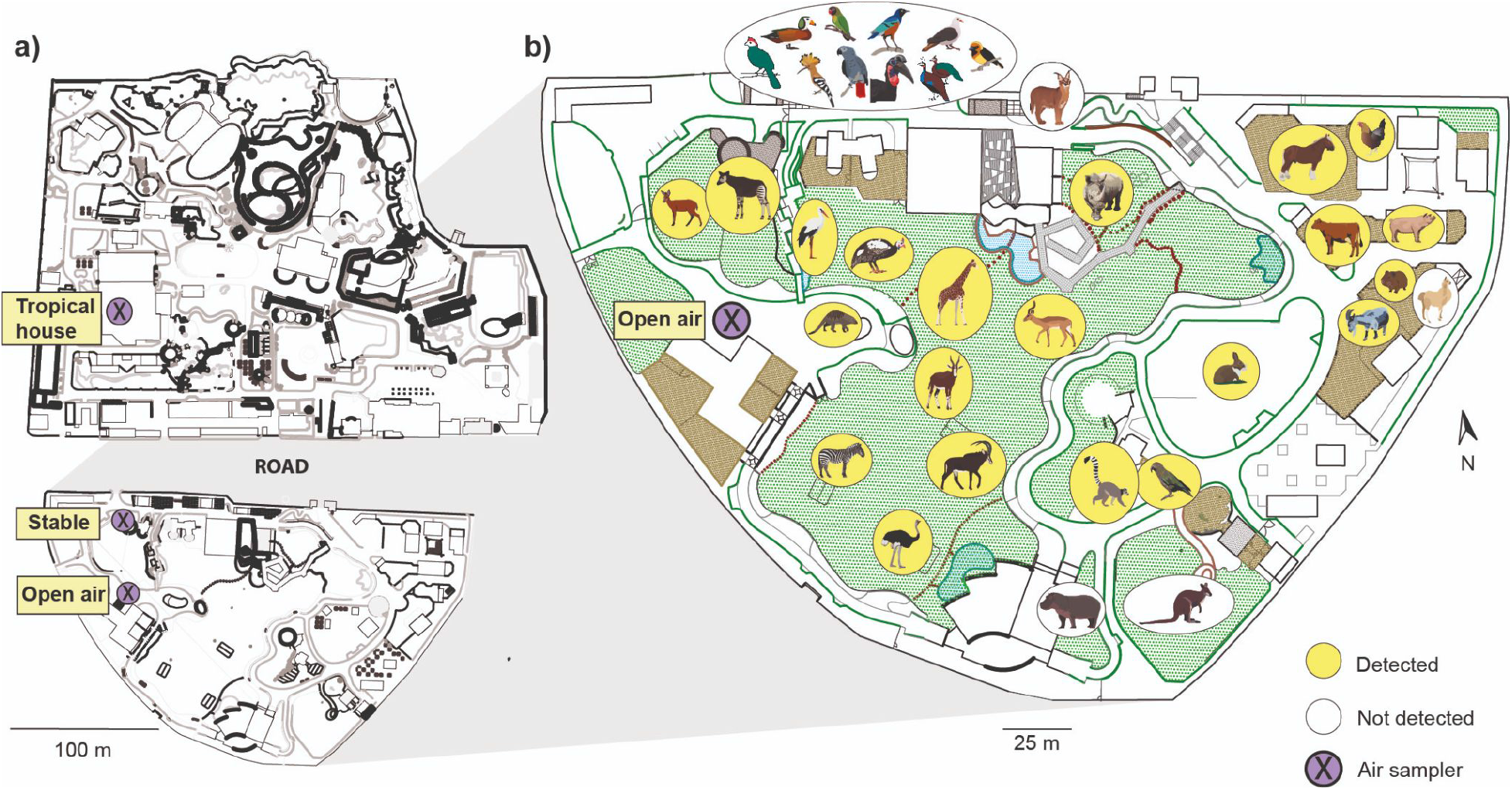
The sampling sites and airDNA detections of vertebrate species. a) The three locations where airDNA samples were collected in Copenhagen Zoo, Denmark: the okapi and red forest duiker stable, in open air among the outdoor enclosures and inside the tropical house. b) AirDNA sampling in open air. Visualised vertebrates have access to outdoor enclosures in the southern part of the zoo. Vertebrate species detected through DNA metabarcoding of airDNA are highlighted in yellow. Maps and animal illustrations courtesy of Copenhagen Zoo.

**Figure 2.**
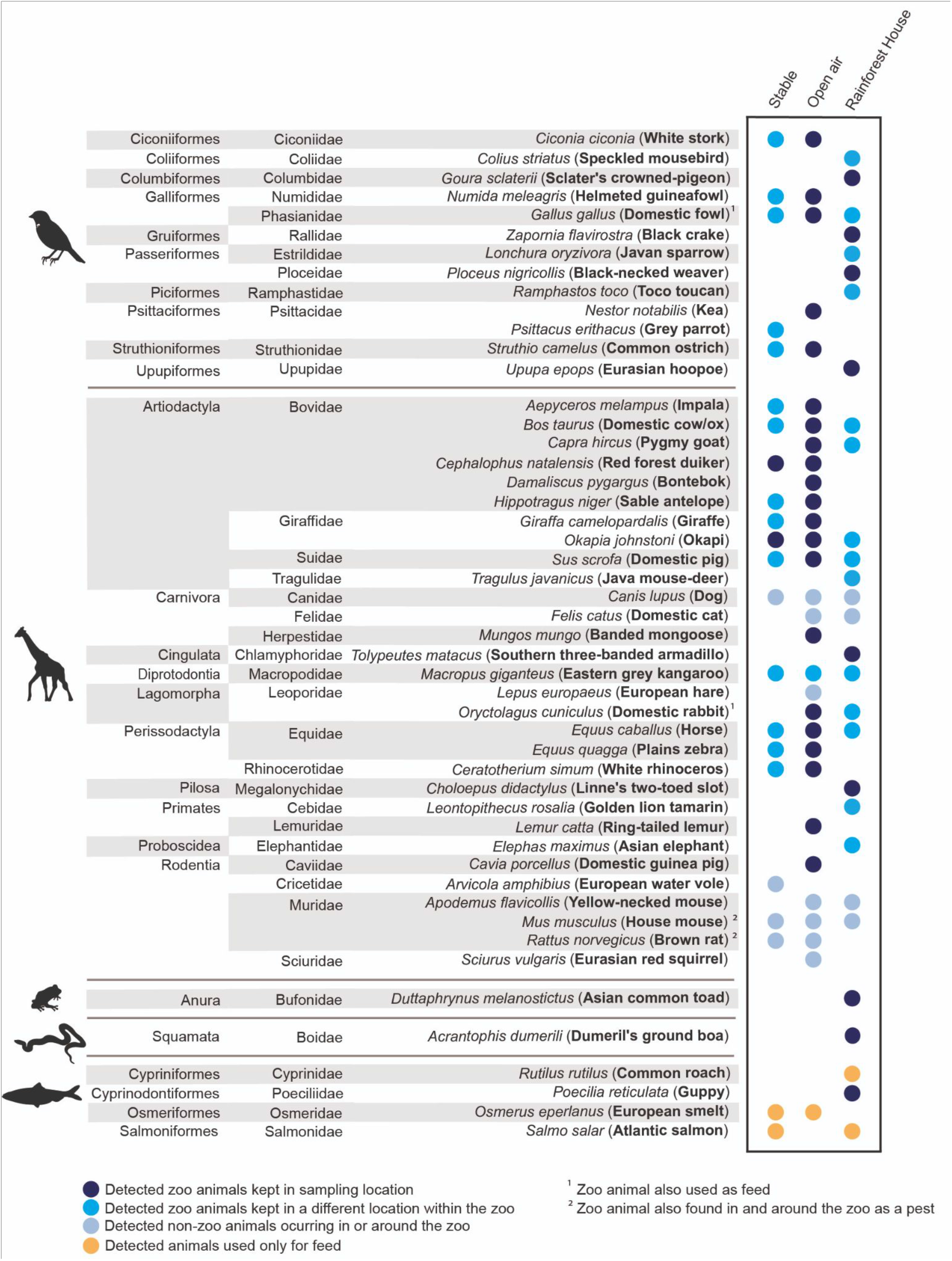
Vertebrate species detected through metabarcoding of airDNA. Detections are made through DNA metabarcoding of 40 samples of airborne particles from three sampling locations in Copenhagen Zoo, Denmark: the okapi and red forest duiker stable (n=12), outside among the outdoor animal enclosures (n=14) and inside the Rainforest House within the Tropical House (n=12). Only taxa that could be determined to species level are included. Taxonomic order and family are listed for each species; common names are in bold. Detected species fall within four categories; detected through air DNA sampling where they are kept (dark blue), detected in another sampling location than where they are kept (blue), detection of wild or domestic non-zoo species (light blue), and species used as animal feed (orange). Some animals kept at the zoo (domestic rabbit and fowl) were also used for feed (1). Further, other animals kept at the zoo (house mouse and brown rat) are known to occur as pests in and around the zoo (2). Detections were made with DNA metabarcoding with two mitochondrial primer sets, one targeting a mammal and one targeting a vertebrate marker. Animal illustrations obtained from the Integration and Application Network (ian.umces.edu/media-library).

To further explore the potential of airDNA to monitor terrestrial vertebrate communities, we deployed air samplers at a location proximal to multiple outdoor mammal and bird enclosures in the southern section of the zoo (Fig. 1b). In total, 16 samples of airborne particulate matter were collected, split between the water vacuum (n = 4 in September; n=4 in December), and the 5 V (n=2 in December) and 24 V samplers (n=6 in December). Between 8 and 21 non-human vertebrates (mean = 14.5, SD= 4.69) were detected in each of the 16 samples (Supplementary Table 2), totalling 30 non-human vertebrate species for the outdoor sampling site (Fig. 2; Supplementary Table 3). Among these, we detected 21 of the 35 bird and mammal species that had access to an outdoor enclosure in the southern section of the zoo (Fig. 1b; Fig. 2). We further detected 1 zoo animal present in the north section of the zoo, 2 animals kept in the zoo but that are also known to be pests (i.e. house mouse and brown rat), 5 wild or domestic non-zoo mammal species known to occur in and around the zoo (e.g. cat and squirrel) and 1 fish species used as feed.

To test whether sequencing of airborne particulate matter would allow detection of taxonomic groups other than birds and mammals, we collected 12 samples inside the Tropical House (Fig. 1a). The Tropical House consists of two main parts, the Butterfly House and the Rainforest House. We sampled in the latter, which contains multiple reptile, bird, and mammal species not present in the outdoor enclosures, except for the Eurasian hoopoe (*Upupa epops*). In the 12 samples collected in the Rainforest House, we detected 7 to 17 non-human vertebrate species per sample (mean = 12.17, SD = 2.98) summing to a total of 29 species, including 16 mammal, 8 bird, 3 fish, 1 amphibian and 1 reptile species (Fig. 2; Supplementary Table 4; Supplementary Table 2). These 29 species included 9 of the 24 vertebrate species kept in the Rainforest House of which 1 of the detected species is kept within a terrarium, namely the Dumeril’s ground boa (*Acrantophis dumerili*). In addition, we detected 5 species kept in other parts of the Tropical House, 4 species used as feed in the zoo, and 7 zoo species kept outside the Tropical House. Further, we detected 2 wild or domestic non-zoo species known to occur in and around the zoo, and 2 rodents known to be pests (Fig. 2). For the total list of species present in the entire Tropical House, see Supplementary Table 5.

We collated all data across sites and samples in an overall inventory. We detected between 9 and 23 non-human vertebrate species per sample (mean = 15.6, SD = 4.06), summing to a total of 49 vertebrate species spanning 26 taxonomic orders and 37 families; 30 mammal, 13 bird, 4 fish, 1 amphibian and 1 reptile species (Fig. 2). Of these 49 species, 38 were exotic animals kept in the zoo, 3 were fish species routinely used as animal feed in the zoo, 2 rodent species kept at the zoo but also known to be pests, and the remaining 6 were wild or domestic non-zoo species known to occur in or around the zoo. Thus, the presence of all 49 detected species could be accounted for. The robustness of our method is further demonstrated by 39 matching species detections between the two sets of sampling replicates, with the remaining 10 taxa only being detected by one of the two sampling replicate sets. These are conservative identifications as they only include those OTU sequences that could be identified to species level. However, for OTU sequences that we could only assign to higher taxonomic levels, we detected Columbidae, a bird family consisting of pigeons and doves, Passeriformes, a large song-bird family, and *Corvus* sp, corvids. These taxonomic groups include wild or feral birds such as jackdaws, crows, pigeons and house sparrows, which are common in and around the zoo.

The detected vertebrates represent species with a large variation in sizes, behaviours and the number of animals present in the zoo, illustrating that a wide range of species can be detected by airDNA sampling. For example, among the species we detected, the zoo holds 2 ostriches (*Struthio camelus*) each weighing ca. 90 kg, 5 white rhinoceros (*Ceratotherium simum*) each weighing ca. 1800 kg, 25 helmeted guineafowls (*Numida meleagris*) each weighing ca. 1.3 kg, and 47 Javan sparrows (*Lonchura oryzivora*) each weighing ca. 22 g. Furthermore, although most of the detected vertebrate species were cursorial (e.g. the impala, *Aepyceros melampus*; and the Java mouse-deer, *Tragulus javanicus*), other lifestyles were also detected, including volant birds (e.g. kea, *Nestor notabilis*), a crawling snake (Dumeril’s ground boa, *Acrantophis dumerili*) and arboreal animals (e.g. two-toed sloth, *Choloepus didactylus*).

### Biomass and distance to air sampling device influence detection

In studies of natural systems, airDNA will predominantly be collected in open air. Thus, we explored putative factors influencing the detection of vertebrate DNA in the outdoor sampling site. This included comparing average biomass and distance from sampler for the species we detected versus those not detected. In addition, we used a logistic regression model with air filtering method, sampling time, average distance of animal to the samplers, animal biomass (no. individuals x average weight for individuals, log transformed) and the taxonomic group (mammal and bird) as independent variables. We found that higher animal biomass (p-value < 0.001) (Fig. 3b) and a shorter distance to the sampler (p-value < 0.05) (Fig. 3c) and significantly increased the probability of vertebrate DNA detection, but found no significant effect of the taxonomic class, the choice of sampling device or sampling time. We hypothesise that larger animals shed more DNA and are more easily detected, per individual. However, when excluding biomass from the model, mammals had a higher probability of being detected than birds (p-value < 0.001).

**Fig. 3.**
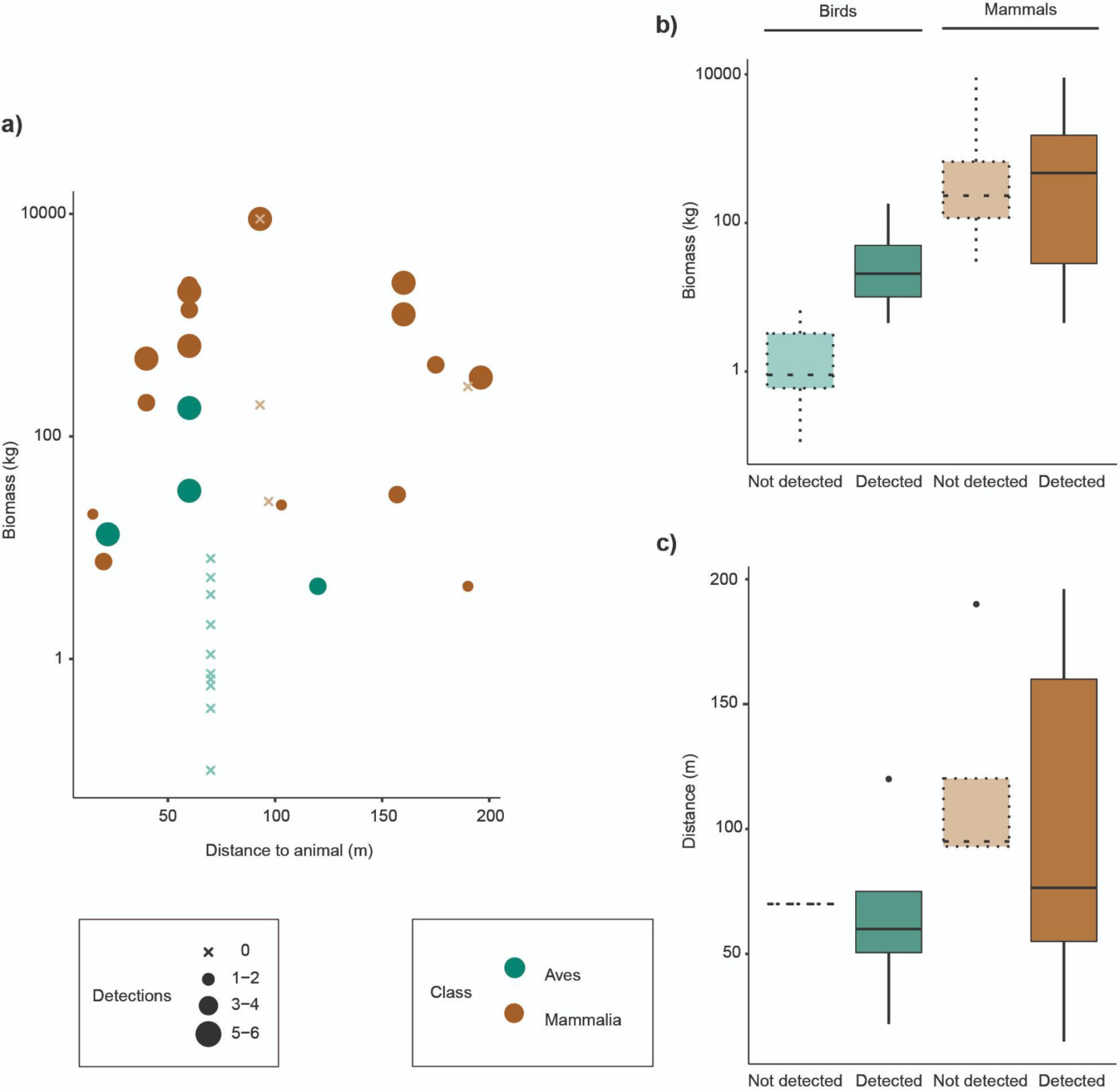
Factors influencing airDNA detections of vertebrate species in open air. The effect of biomass, distance on species detections. Only data from zoo animals with access to an outdoor enclosure in the southern part of the zoo, whether they were detected or not. a) Influence of biomass and distance on the number of times a species was detected across the six different sampling events (i.e. water vacuum for 30 and 60 min, particle filter sampler with the 24 V sampler ran during 30, 60 and 300 min, and the 5 V sampler ran during 30 hrs). b) Average biomass estimated as weight times the number of animals for the bird and mammal species that were detected and not detected in any of the air samples. c) Average distance between the samplers and open air enclosures for the bird and mammal species that were detected and not detected in any of the air samples.

The concentration of DNA is expected to fall with distance from the source, and accordingly we only detected one of the species located in the northern section of the zoo, namely the eastern grey kangaroo (*Macropus giganteus*). We speculate that distance and the presence of several buildings and a trafficked road between our sampling site in the southern part of the zoo and the northern part prevented us from detecting other vertebrate species present in the northern part of the zoo. Similarly, the failure to detect 14 species present in the southern part of the zoo could not only be due to the biomass, distance and taxonomic groups, but also the presence of buildings between the enclosure and the air sampler (Fig. 1b

Despite not finding significant differences in detections between samplers, we did observe practical differences. We found the water vacuum sampler to be more noisy and less flexible due to its size and the need for an external power supply and molecular grade sterilized water. In contrast, the two particle samplers have the advantage of being small and portable, which allows them to be deployed in a wide variety of environments for several days depending on the power supply. The versatility comes at the expense of airflow, as the particle filter sampler with the 24 V blower fan provides a larger airflow of about 0.8 m^3^/min compared to the particle filter sampler with the 5 V blower fan with about 0.03 m^3^/min. Nevertheless, the compact size of both samplers and their very low noise level (the 24 VDC blower fan is rated at 55 dB-A) makes them suitable for environments where wildlife is easily disturbed. As it can be assumed that wildlife-DNA will often travel in association with airborne dust and fibers, typically in the size range of 1 μm-10 μm, using a less dense filter than the F8 used herein, would both decrease the collection efficiency towards smaller particles and increase the airflow through the filter. It is therefore necessary to consider the product of the two (collection efficiency * airflow) when calculating the effective sampling volume.

### Implications for monitoring of terrestrial vertebrate communities

Our results suggest that airDNA is an untapped source of spatial and temporal vertebrate distribution data with the potential to transform the way natural ecosystems are studied and surveyed. This includes acting as a cost-effective and efficient tool to inform conservation efforts, evaluate sustainable removal levels, and track progress in achieving biodiversity targets, something of great global importance given the ongoing climate and biodiversity crisis ^7–9^.

We carried out the study at the Copenhagen Zoo which provided a suitable controlled source due to the presence of well-defined individual animals. However, their confinement and density in the enclosure may have artificially increased their probability of being detected in air samples compared to sampling in a natural environment. Still, we detected six non-zoo animals in the air samples despite the high zoo species biomass and concentration as compared to non-zoo animals in the surrounding area.

As with any novel methodology, including the first demonstrations of eDNA in aquatic environments ^15,54,55^, the full potential of airDNA for vertebrate community surveys will require further optimisations and developments across a range of natural habitats and applications before standardisation and implementation in routine monitoring can be achieved. With time, we envision terrestrial airDNA vertebrate surveys could parallel the field of aquatic eDNA monitoring with the potential to revolutionize and form the cornerstone in future ecosystem studies, including global next generation biomonitoring frameworks ^4,5^

## Author contributions

CL, MTO and KB conceived and designed the study with input from CVJ, MFB and MSJ; MTO and KB provided funding; CL, CVJ and MSJ designed and tested the particle samplers; CL performed the sampling and the lab work; CL, MFB, TGF, MTO and KB analysed the data; CL, CVJ, MTO and KB drafted the paper. All authors read and approved of the final version.

## Acknowledgments

This study was funded by an Experiment grant (grant no. 00028049) from VILLUM FONDEN to MTO and KB. We thank Ruta Sidaraviciute from Airlabs for helping with the design and 3D-printing of the particle samplers. We thank the Copenhagen Zoo staff for facilitating the sample collection and Frederik Leerhøi, Physilia Chua, Åshild Vågene, Sarah Mak, Jonas Niemann, and Camilla Pereira for assistance with sample collection. We thank Kurt Kjær, Fabian Roger, Lasse Vinner and Mette Juul Jacobsen for discussions, and Hannes Schoeder and Tom Gilbert for comments on the original manuscript.

## Methods

### Study site

Fortyfour air samples were collected at the Copenhagen Zoo, Denmark, during September and December 2020. Air samples were collected in three places: 1) inside a 155 m^2^ stable inhabited by two okapis (*Okapi johnstoni*) and two red duiker (*Cephalophus natalensis*), which had the option to also use an adjoining outdoor enclosure during the day, 2) we sampled outdoors in open air at a fixed location in the part of the Zoo containing multiple outdoor enclosures with a mixed variety of mammals and other terrestrial vertebrates. Finally, 3) inside the Rainforest House, a 442 m^2^/2200 m^3^ confined enclosure in which smaller vertebrates and other animals from the tropics move freely around both day and night located within the Tropical House (Fig 1; Supplementary Table 5). Inside the stable and the Rainforest House, the temperature was kept constant during September and December, ranging from 18.6 to 20.5°C, and from 22 to 27°C, respectively. None of the locations were directly exposed to the wind, but the stable did have openings to the outdoors and Rainforest House had an internal ventilation system. During the outdoor sample collection, on the 11th of September the temperature ranged from 17.1 to 17.3°C, wind speed from 4.4 to 5.2 m/s with wind coming from the SW; on the 22th of September the temperature ranged from 21 to 21.2 °C, wind speed of 2.6 to 4.5 m/s and with wind coming from S; on the 10th of December, the temperature was 3°C, wind speed was 5.9 to 5 m/s and the wind came from the East; and on the 11th of December, the temperature was 2.4°C, wind speed 5.6 to 5.4 m/s and the wind came from SE/E direction (dmi.dk).

### AirDNA samplers

Environmental DNA was collected from air using three different samplers, a water-based commercial vacuum and two air particle filters with different power sources and airflows. The water-based commercial vacuum was the Kärcher DS5800 Water Vacuum (WV) (Alfred Kärcher GmbH & Co. KG, Germany), which consists of a high-flow-rate impinger with an outer part that creates suction and an inner vortex chamber where the particles flow into ^56^. This WV was connected to the electrical circuit and provided an average airflow of 8.8 m^3^/min.

The second sampler was a custom made air particle filter sampler consisting of a Delta Electronics 97.2 mm x 33 mm 24 V, 0.550 A DC brushless radial blower fan, a class F8 pleated fibrous particulate filter (Dongguan Wonen Environmental Protection Technology Co.,Ltd), and a 3D-printed filter housing (Airlabs, Copenhagen, Denmark; 3D-printing blueprints available in Supplementary Information). The filter was placed approximately 40 mm from the intake of the blower fan and was connected to the electrical circuit, providing an airflow of 0.8 m^3^/min. We call this the medium filter (MF) sampler.

The third sampler was overall similar to the MF sampler, except that the filter was placed on a 3D-printed hilter housing approximately 20 mm from the intake of the blower fan, which is a battery-driven Hawkung/Long Sheng Xin 40 mm x 40 mm x 10 mm, 5 V, 0.10 A DC brushless radial blower fan, providing an airflow of 0.03 m^3^/min (3D-printing blueprints available in Supplementary Information). We call this the small filter (SF) sampler. For both MF and SF samplers, we used class F8 pleated fibrous particulate filters (Dongguan Wonen Environmental Protection Technology Co.,Ltd). This type of filter is usually implemented in A/C units and is designed to capture airborne particulate matter and micro- and nanofibers with high efficiency and low pressure drop. As the filter is cut and stretched out to a single layer around the size of the filter housing, the airflow and retention efficiency is expected to decrease slightly from the official F8 rating (https://www.emw.de/en/filter-campus/filter-classes.html).

### Sample collection

Sampling with the WV sampler followed ^56^, i.e. the inner vortex chamber was filled with 1.7 L sterile Milli-Q H2O. After running the impinger, the water from the vortex chamber was filtered using Sterivex filters (pore size 0.22 μm). In between samplings, the vortex chamber and the suction hole were cleaned with 5% sodium hypochlorite (bleach) and 70% ethanol. At every location, a sampling negative control consisting of 200 mL of sterile Milli-Q H2O was added to the vortex chamber and thereafter filtered with Sterivex filters. Using this sampler, air was collected for 30 min and 60 min at each site during December. Samples collected outside with the WV were also collected during September. Prior to sampling with the MF and SF air filter samplers, the F8 filters were cut into a smaller size to fit the housing, autoclaved, placed under UV light for 20 min and thereafter stored individually in sterile plastic bags. In between sampling events, the housing of the MF and SF samplers was cleaned with 5% bleach and 70% ethanol. Sterilized filters were handled using sterile tweezers and stored in a sterile 50 mL Falcon tube upon sampling. To test the effect of sampling time, the MF sampler was left running for 30 min, 60 min and 5 hrs. For the SF sampler, to test the effect of long sampling time, this sampler was left running for 30 hrs at each location. Both samplers were run during December. For all three samplers, samples were taken at 1 m above the ground and in duplicates. Filters were stored in a cooling box for up to 5 hours and thereafter at −20 °C until DNA extraction.

### DNA extraction

Due to their big size, the MF filters were cut in half with sterile blades. Both halves of MF filters and entire SF filters were transferred independently to 5 mL Eppendorf tubes and 3 mL of autoclaved PBS pH 7.4 (1X) (Gibco™, Thermo Fisher) was added. After an incubation of 45 mins, the filters were transferred to a new Eppendorf tube, and the PBS was centrifuged at 6000 xg during 10 min to create a pellet, and the supernatant removed. The DNeasy Blood & Tissue Kit (Qiagen, USA) was used for DNA extraction of the PBS pellets, the two halves of the MF filters, the entire SF filters and the Sterivex filters from the VW sampler. In addition, to test for contamination in the sterilized MF and SF filters, Sterivex filters and the autoclaved PBS, non-used filters and PBS were subjected to DNA extraction. In addition, to test for contamination in the DNA extraction room, two falcon tubes containing 50 mL sterile Milli-Q H2O left open for 48 hrs, and were also subjected to DNA extraction.

The DNA extraction followed manufacturer’s instructions (Purification of Total DNA from Animal Tissues protocol), with slight modifications: the ratio 9:1 of ATL buffer to Proteinase K was kept but the volume was increased to 720 ATL and 80 Proteinase K and an incubation step of 37 °C for 15 min was added after the addition of 50 μl of EBT (EB buffer with 0.05% Tween-20 (VWR)). This elution step was carried out twice to increase DNA yield.

As belonging to the same sample, the digest of the filters and from the PBS pellet were passed through the same spin column and therefore having one DNA extract per sample. However, for three MF samples collected inside the Okapi stable, the digests of each half of the filter presented many particles clogging the spin column and therefore the digests could not be combined into one spin column. This resulted in a total of 49 DNA extracts, representing 40 samples (see Supplementary Table 7). Negative extraction controls were added for every 16 samples. Eluted DNA was stored in Eppendorf LoBind tubes at −20°C.

To minimize contamination risk during DNA extraction, we set up a specialised environmental DNA pre-PCR laboratory, which was thoroughly cleaned prior to its use and in which guidelines follow those used in ancient DNA laboratories such as the use of hair net, sleeves, facemask, two layers of medical gloves, dedicated footwear and the use of ≥3% bleach on all surfaces ^57^.

## DNA metabarcoding

Metabarcoding was conducted using two different primer sets. To target mammals, a ca. 95 bp 16S rRNA mitochondrial marker was PCR-amplified with the primers 16Smam1 (forward 5’-CGGTTGGGGTGACCTCGGA-3’) and 16Smam2 (reverse 5’-GCTGTTATCCCTAGGGTAACT-3’) (Taylor, 1996). To target vertebrates, a ca. 97 bp fragment of the 12S gene was PCR-amplified with the primer set 12SV05 forward 5’-TTAGATACCCCACTATGC-3’ and 12SV05 reverse 5’-TAGAACAGGCTCCTCTAG-3’ (Riaz et al., 2011). The two metabarcoding primer sets are from here on referred to as 16S mammal and 12S vertebrate primers, respectively. Nucleotide tags were added to the 5’ ends of forward and reverse primers of both primer sets to allow parallel sequencing ^58^. Nucleotide tags were six nucleotide tags in length and had min. 3 nucleotide differences between them. One to two nucleotides were added to the 5’end to increase complexity on the flowcell. DNA extracts from fin whale (*Balaenoptera physalus*) and bowhead whale (*Balaena mysticetus*) were used as positive controls, as none of the species are found close to the sampling site in Copenhagen Zoo.

Prior to metabarcoding PCR amplification, dilution series of a subset of the DNA extracts were screened using SYBR Green quantitative PCR (qPCR). This was done to determine the optimal cycle number and DNA template volume to ensure optimal amplification in the following metabarcoding PCR amplifications. Further, all negative controls were included in the qPCR to screen for contamination.

For the 16S mammal primer, the 20 μl reactions consisted of 2 or 4 μl DNA template, 0.75 U AmpliTaq Gold, 1× Gold PCR Buffer, and 2.5 mM MgCl2 (all from Applied Biosystems); 0.6 μM each of 5’ nucleotide tagged forward and reverse primer; 0.2 mM dNTP mix (Invitrogen); 0.5 mg/ml bovine serum albumin (BSA, Bio Labs); 3 μM human blocker (5’– 3’ GCGACCTCGGAGCAGAACCC–spacerC3) ^59^; and 1μL of SYBR Green/ROX solution [one part SYBR Green I nucleic acid gel stain (S7563) (Invitrogen), four parts ROX Reference Dye (12223-012) (Invitrogen) and 2000 parts high-grade DMSO]. The thermal cycling profile was 95°C for 10 min, followed by 40 cycles of 95°C for 12 s, 59°C for 30 s, and 70°C for 25 s, followed by a dissociation curve. For the 12SVert primer, the 20 μl reaction was the same except for the human blocker (5’–3’ TACCCCACTATGCTTAGCCCTAAACCTCAACAGTTAAATC– spacerC3) ^32^ and the thermal cycling profile of 95°C for 10 min, followed by 40 cycles of 94°C for 30 s, 59°C for 45 s, and 72°C for 60 s, followed by a dissociation curve. The amplification plots from the qPCR indicated that 2 μl DNA template was optimal, 35 and 38 cycles were optimal for the 16S mammal and 12S vertebrate primers, respectively, and the negative extraction controls showed no contamination.

For the metabarcoding PCR, the 20 μL reactions were set up as described for the qPCR above but omitting SYBR Green/ROX and replacing the dissociation curve with a final extension time of 72°C for 7 min. Four tagged PCR replicates were carried out for each of the 49 DNA extracts, negative and positive controls, and for both primer sets; PCR replicates from each sample were differently tagged. Negative controls were included every seven PCR reactions.

Amplified PCR products were visualized on 2% agarose gels with GelRed against a 50 bp ladder. All negative controls appeared negative and all positive controls showed successful amplification. Even if not showing a successful amplification, all PCR products of DNA extracts, including negative and positive controls carrying different nucleotide tag combinations, were pooled resulting in four amplicon pools: one pool per replicate.

Amplicon pools were purified with MagBio HiPrep beads (LabLife) using a 1.6x bead to amplicon pool ratio and eluted in 35 μL EB buffer (Qiagen). Purified amplicon pools were built into sequence libraries with the TagSteady protocol to avoid tag-jumping (Carøe & Bohmann, 2020). Libraries were purified with a 1.6x bead to library ratio and eluted in 30 μL EB buffer and qPCR quantified using the NEBNext Library Quant Kit for Illumina (New England BioLabs Inc.). Purified libraries were pooled equimolarly according to the qPCR results and sequenced at the GeoGenetics Sequencing Core, University of Copenhagen, Denmark. Libraries were sequenced using 150 bp paired-end reads on an Illumina MiSeq sequencing platform using v3 chemistry aiming at 30,000 reads per PCR replicate.

## Data analysis

Sequence data for each primer set was processed separately. Illumina adapters and low quality reads were removed and paired ends merged using AdapterRemoval v2.2.2^60^. Within each amplicon library, sequences were sorted based on primers and tag sequences using Begum ^61^ allowing two primer-to-sequence mismatches. Further, for each sample Begum was used to filter sequences across the PCR replicates guided by the positive and negative controls and retaining sequences found in three out of the four PCR replicates and with a minimum copy number of 10 and 6 for the 16S mammal and 12S vertebrate primer sets, respectively. As the aim of the present study was to detect and identify species and not intraspecific variation, we decided to create clusters of sequences, instead of denoising and creating amplicon sequence variants (ASV). Clustering and denoising have proven to be complementary, instead of alternatives, and when working with eukaryotes, clustering should be the standard unit as long as using the correct parameter settings during data analysis ^62^. The filtered sequences with a similarity score of 97% were therefore clustered into operational taxonomic units (OTU) using SUMACLUST ^63^. Curation of the OTUs was carried out with the LULU algorithm ^64^, using default settings to remove erroneous OTUs.

Taxonomic identification of the OTU sequences was carried out using BLASTn against the NCBI Genbank database. The output was imported into MEGAN Community Edition v6.12.7 ^65^ using a weighted LCA algorithm with 80% coverage, top percent of 10, and a minimum score of 150. The taxonomic identification of all OTU sequences was manually checked to validate them and species-level identification was assigned if the OTU sequence had a 100% identity match to a NCBI reference sequence. We assigned those that matched 100% to more than one species to the species found in the Copenhagen Zoo. In a few cases where multiple OTUs were assigned to the same species, the corresponding DNA sequences were checked visually in Geneious Prime 2020.1.2 to assess whether the OTUs resulted from genuine haplotype variation or biases caused by minor variations in sequence length. OTUs that could not be identified to species level were discarded before further analysis. In addition, sequences matching to human were removed, as well as those matching to chimpanzee (*Pan troglodytes*) due to its close similarity to human sequences. DNA from the Sclater’s crowned pigeon (*Goura sclateri*) was detected in the water vacuum samples collected at the open-air location, but as it was also detected in the sampling negative control, it was considered cross-contamination and therefore deleted from the data from that site. One of the few non-detected mammals in the outdoor sampling was wallaby, whereas the Eastern grey kangaroo was detected, even though it is found on the North part of the zoo. Both animals belong to the same genus, *Macropus*, but both markers show a 100% match to kangaroo DNA. Finally, the OTU taxonomically identified as *Canis lupus* could originate from dog or grey wolf. Although three grey wolves were present in the zoo during the sampling in September, they were absent during the sampling in December. Further, the detection of this OTU in all samplers made us conclude that the DNA detected is from dogs in the area. We collated data across replicates and the two primer sets in an overall inventory.

For the statistical analysis only data from species present at the southern part of the zoo was used. Detected noon-zoo animals and those also used as feed were removed from the dataset, as it was not possible to measure the exact location and biomass. The distance of the animals to the samplers was measured using a satellite view of the Copenhagen Zoo using Google earth (https://earth.google.com/) and using an average point of reference for animals with a large enclosure. Average body weight data was obtained from Species360 Zoological Information Management System (ZIMS) (2021). The dataset used can be found in Supplementary Table 6. We fitted a logistic model (estimated using ML) to predict detection with distance, biomass (log transformed), taxonomic group (class: bird or mammal), sampler type (WV, SF, MF) and sampling time as potential explanatory variables. Effect of sampler type, taxonomic group and sampling time were insignificant and they were removed from the model. The explanatory power of the final model (formula: detection ~ distance + log(biomass)) was substantial (Tjur’s R^2^ = 0.30). The model’s intercept, corresponding to distance = 0 and biomass = 0, is at −1.50 (95% CI [−2.29, −0.76], p < .001). Within this model: the effect of distance is statistically significant and negative (beta =−8.04e-03, 95% CI [−0.01, −1.68e-03], p = 0.015; Std. beta = −0.14, 95% CI [−0.44, 0.15]); the effect of biomass(kg) (log transformed) is statistically significant and positive (beta = 0.47, 95% CI [0.34, 0.61], p < .001; Std. beta = 1.57, 95% CI [0.78, 2.52]). When not using animal biomass as a potential explanatory variable in the model, the effect of taxonomic group was significant. In this case, the explanatory power of this model (formula: value ~ Distance + Class) was weak (Tjur’s R^2^ = 0.10). The model’s intercept, corresponding to distance = 0 and Class = Aves, is at −0.67 (95% CI [−1.33, −0.04], p = 0.040). Within this model, the effect of distance is statistically significant and negative (beta = −6.22e-03, 95% CI [−0.01, −1.88e-04], p = 0.046; Std. beta = −0.31, 95% CI [−0.62, −9.31e-03]). The effect of Class [Mammalia] is statistically significant and positive (beta = 1.45, 95% CI [0.81, 2.12], p < .001; Std. beta = 1.45, 95% CI [0.81, 2.12]). The data used for the logistic model can be found in Supplementary Table 6.

## Authenticity

Metabarcoding with universal primers that PCR amplify short fragments of the often low DNA quantities of target taxa in environmental DNA extracts comes with the inherent risk of amplifying contaminant templates ^e.g. 66^. Further, library preparation, PCR and sequencing artefacts can lead to inflated diversity and false positives ^67^. To ensure authenticity of results, we therefore followed strict sampling, laboratoratory and bioinformatic workflows.

To minimise risk of contamination, we created a dedicated specialised environmental DNA pre-PCR laboratory for DNA extractions of the air filtering samples in which we set up and followed guidelines commonly used for ancient DNA laboratories, such as unidirectional workflow and the use of hair net, sleeves, facemask, two layers of medical gloves, dedicated footwear, decontamination with ≥3% bleach^57^. All steps of the workflow were carried out in laminar flowhoods and using filter tips. To reduce risk of PCR introduced artefacts, we carried out only one PCR amplification for each sample PCR replicate prior to sequencing. To ensure that tag-jumps or index switching would not cause spillover in samples, we used a library preparation protocol that allowed avoidance of tag-jumps, i.e. false assignment of sequences to samples ^68^, and twin dual-indexes to ensure that potential library bleeding would not cause false assignment of sequences to libraries ^69,70^.

To enable identification of potential contamination, we included negative controls in all steps of the workflow and positive controls in the metabarcoding PCR amplifications. During laboratory quality control steps, we did not identify contamination in any of the negative controls. Despite this, we sequenced the negative and positive controls alongside the samples. The positive control species were not found in the sampling area, which allowed us to assess spillover between samples. We did not detect any spillover from the positive controls to the samples or negative controls, which indicated that there was no cross-contamination in the metabarcoding PCR and the following downstream analyses. No negative controls contained OTUs, except for one of the negative sampling control from the water vacuum. Here, an OTU from a bird only found inside the Rainforest House (Sclater’s crowned-pigeon) was detected. This OTU was further detected in the water vacuum samples collected outdoors, which took place after the collection in the Rainforest House. We detected no other OTUs from the Rainforest House in the water vacuum samples collected outside. We therefore excluded this cross-contaminating OTU from all water vacuum sampler detections. The other taxa detected with this sampler were not detected in the negative sampling controls, and were therefore determined to be true detections. No vertebrate DNA was detected in the negative controls from the 24 V and 5 V particle filters.

A crucial step to ensure authenticity was the inclusion of four PCR replicates for all samples and negative and positive controls. This was done for both markers. For each set of four PCR replicates, we used different tag combinations to lower risk of primer cross-contamination and importantly, to allow stringent filtering of sequences across each sample’s PCR replicates ^67,71,72^. We employed a conservative approach in which we only retained sequences that were found in min. three of the four PCR replicates from each sample ^67^. Further, we only retained sequences present in a certain copy number threshold. These were crucial steps that allowed us to balance error removal with detection of diversity ^67^. In addition, we used the LULU algorithm which removes artefactual OTUs and thereby reduces the number of false positives ^64^.

The study was carried out in a zoo environment, which enabled us to verify the presence of all detected taxa in our collated species inventory. We are aware that we detected species known to be contaminants in laboratory reagents, such as pig, cow and chicken and to a lesser extent mouse, rabbit, goat and guinea pig ^73^. For example, we used bovine serum albumin (BSA) in PCR amplifications which is synthesised from cow’s blood ^74^. We did however not detect any of these taxa in the negative controls. Further, cow, pig, chicken, mouse, rabbit, goat and guinea pig are present in the zoo and we therefore find our detections of them in air particle samples reliable.

## Websites

IUCN 2021. The IUCN Red List of Threatened Species. Version 2021-1. https://www.iucnredlist.org. Downloaded 28 June 2021.

Danish Meteorological Institute, Weather archive. https://www.dmi.dk/vejrarkiv/ (2020). Species360 Zoological Information Management System (ZIMS) (2021). zims.Species360.org

