## Supplementary Information for "Airborne environmental DNA for terrestrial vertebrate community monitoring"

### Supplementary tables:

**Supplementary Table 1.** List of non-human vertebrate species detected in air samples (n=12) collected in the okapi and red forest duiker stable, by using the 16S mammal (m) and the 12S vertebrate (v) metabarcoding primers. Detections range from non-detected (0) to detection in both replicates (2) in the different samplers used: using water vacuum (WV) for 30 and 60 min, particle filter sampler with the 24V fan (MF) ran during 30, 60 and 5hrs, and particle filter sampler with the 5V fan (SF) ran during 30 hrs. Information regarding the presence of the species within the stable, and in the rest of the zoo, is provided.

| Class | Order | Family | Species | Common name | Primer | WV_30min | WV_60min | MF_30min | MF_60min | MF_5hrs | SF_30hrs | Zoo animal | Present in stable |
|---|---|---|---|---|---|---|---|---|---|---|---|---|---|
| Actinopterygii | Clupeiformes | Clupeidae | NA | NA | v | 0 | 0 | 0 | 0 | 1 | 0 | Food | No |
| | Osmeriformes | Osmeridae | *Osmerus eperlanus* | European smelt | v | 0 | 0 | 0 | 0 | 1 | 0 | Food | No |
| | Salmoniformes | Salmonidae | *Salmo salar* | Atlantic salmon | v | 0 | 1 | 0 | 1 | 0 | 0 | Food | No |
| Aves | Anseriformes | Anatidae | NA | NA | v | 0 | 0 | 1 | 0 | 2 | 0 | NA | NA |
| | Ciconiiformes | Ciconiidae | *Ciconia ciconia* | White stork | v | 0 | 2 | 2 | 1 | 1 | 2 | Yes | No |
| | Columbiformes | Columbidae | NA | NA | v | 2 | 2 | 2 | 2 | 2 | 2 | Yes | No |
| | Galliformes | Numididae | *Numida meleagris* | Helmeted guineafowl | v | 1 | 2 | 2 | 2 | 2 | 2 | Yes | No |
| | | Phasianidae | *Gallus gallus* | Domestic fowl | v | 2 | 2 | 2 | 0 | 2 | 2 | Yes | No |
| | Passeriformes | Corvidae | NA | NA | v | 2 | 2 | 1 | 0 | 2 | 2 | NA | NA |
| | | Ploceidae | *Ploceus* sp. | Weaver | v | 0 | 1 | 0 | 0 | 0 | 0 | Yes | No |
| | | NA | NA | NA | v | 1 | 0 | 0 | 0 | 2 | 0 | NA | NA |
| | Psittaciformes | Psittacidae | *Psittacus erithacus* | Grey parrot | v | 1 | 2 | 0 | 0 | 1 | 0 | Yes | No |
| | Struthioniformes | Struthionidae | *Struthio camelus* | Common ostrich | v | 0 | 1 | 0 | 0 | 2 | 2 | Yes | No |

| | | | | | | | | | | | | | |
|---|---|---|---|---|---|---|---|---|---|---|---|---|---|
| Mammalia | Artiodactyla | Bovidae | *Aepyceros melampus* | Impala | m/v | 1 | 1 | 2 | 0 | 1 | 0 | Yes | No |
| | | | *Bos taurus* | Domestic cow/ox | m/v | 1 | 2 | 1 | 0 | 0 | 0 | Yes | No |
| | | | *Cephalophus natalensis* | Red forest duiker | m | 2 | 2 | 2 | 2 | 2 | 2 | Yes | Yes |
| | | | *Hippotragus niger* | Sable antelope | m/v | 1 | 2 | 1 | 0 | 1 | 2 | Yes | No |
| | | | NA | NA | v | 0 | 2 | 0 | 1 | 2 | 2 | NA | NA |
| | | Giraffidae | *Giraffa camelopardalis* | Giraffe | m/v | 1 | 2 | 2 | 1 | 2 | 2 | Yes | No |
| | | | *Okapia johnstoni* | Okapi | m/v | 2 | 2 | 2 | 2 | 2 | 2 | Yes | Yes |
| | | Suidae | *Sus scrofa* | Domestic pig | m/v | 2 | 2 | 2 | 2 | 2 | 2 | Yes | No |
| | Carnivora | Canidae | *Canis lupus* | Dog / gray wolf | m/v | 2 | 2 | 2 | 2 | 0 | 1 | No | No |
| | Diprotodontia | Macropodidae | *Macropus giganteus* | Eastern grey kangaroo | v | 0 | 0 | 1 | 0 | 1 | 1 | Yes | No |
| | Perissodactyla | Equidae | *Equus caballus* | Horse | m | 0 | 1 | 0 | 0 | 0 | 0 | Yes | No |
| | | | *Equus quagga* | Plains zebra | v | 1 | 0 | 0 | 0 | 0 | 0 | Yes | No |
| | | Rhinocerotidae | *Ceratotherium simum* | White rhinoceros | v | 2 | 2 | 0 | 0 | 2 | 2 | Yes | No |
| | Rodentia | Cricetidae | *Arvicola amphibius* | European water vole | m | 0 | 1 | 0 | 0 | 0 | 0 | No | No |
| | | | NA | NA | m | 0 | 1 | 0 | 0 | 0 | 0 | NA | NA |
| | | Muridae | *Apodemus* sp. | NA | v | 0 | 1 | 2 | 1 | 2 | 2 | No | No |
| | | | *Mus musculus* | House mouse | m/v | 1 | 2 | 2 | 1 | 2 | 2 | Yes | No |
| | | | *Rattus norvegicus* | Brown rat | m | 0 | 1 | 0 | 0 | 0 | 0 | Yes | No |

**Supplementary Table 2.** List of non-human vertebrate species detected in air samples in the three locations within Copenhagen Zoo. Detections are divided per replicate (R1 and R2) for each sample. Non-detected (0) species and detection in each replicates (1) in the different samplers used: using water vacuum (WV) for 30 and 60 min, particle filter sampler with the 24V fan (MF) ran during 30, 60 and 5hrs, and particle filter sampler with the 5V fan(SF) ran during 30 hrs.

| Class | Species | Stable | | | | | | | | | | | | Outdoors | | | | | | | | | | | | | | | | Rainforest House | | | | | | | | | | | |
|---|---|---|---|---|---|---|---|---|---|---|---|---|---|---|---|---|---|---|---|---|---|---|---|---|---|---|---|---|---|---|---|---|---|---|---|---|---|---|---|---|---|
| | | December | | | | | | | | | | | | September | | | | December | | | | | | | | | | | | December | | | | | | | | | | | |
| | | WV_30min | | WV_60min | | MF_30 | | MF_60min | | MF_5hrs | | SF_30hrs | | WV_30min | | WV_60min | | WV_30 | | WV_60min | | MF_30min | | MF_60min | | MF_5hrs | | SF_30hrs | | WV_30min | | WV_60min | | MF_30min | | MF_60min | | MF_5hrs | | SF_30hrs | |
| | | R1 | R2 | R1 | R2 | R1 | R2 | R1 | R2 | R1 | R2 | R1 | R2 | R1 | R2 | R1 | R2 | R1 | R2 | R1 | R2 | R1 | R2 | R1 | R2 | R1 | R2 | R1 | R2 | R1 | R2 | R1 | R2 | R1 | R2 | R1 | R2 | R1 | R2 | R1 | R2 |
| Actinopterygii | *Rutilus rutilus* | 0 | 0 | 0 | 0 | 0 | 0 | 0 | 0 | 0 | 0 | 0 | 0 | 0 | 0 | 0 | 0 | 0 | 0 | 0 | 0 | 0 | 0 | 0 | 0 | 0 | 0 | 0 | 0 | 1 | 0 | 0 | 0 | 0 | 0 | 0 | 0 | 0 | 0 | 0 | 0 |
| | *Poecilia reticulata* | 0 | 0 | 0 | 0 | 0 | 0 | 0 | 0 | 0 | 0 | 0 | 0 | 0 | 0 | 0 | 0 | 0 | 0 | 0 | 0 | 0 | 0 | 0 | 0 | 0 | 0 | 0 | 0 | 1 | 1 | 1 | 1 | 1 | 1 | 0 | 1 | 1 | 0 | 0 | 0 |
| | *Osmerus eperlanus* | 0 | 0 | 0 | 0 | 0 | 0 | 0 | 0 | 1 | 0 | 0 | 0 | 0 | 0 | 1 | 0 | 0 | 0 | 0 | 1 | 0 | 0 | 0 | 0 | 0 | 1 | 0 | 0 | 0 | 0 | 0 | 0 | 0 | 0 | 0 | 0 | 0 | 0 | 0 | 0 |
| | *Salmo salar* | 0 | 0 | 1 | 0 | 0 | 0 | 1 | 0 | 0 | 0 | 0 | 0 | 0 | 0 | 0 | 0 | 0 | 0 | 0 | 0 | 0 | 0 | 0 | 0 | 0 | 0 | 0 | 0 | 0 | 0 | 0 | 1 | 0 | 0 | 0 | 0 | 0 | 0 | 0 | 0 |
| Amphibia | *Duttaphrynus melanostictus* | 0 | 0 | 0 | 0 | 0 | 0 | 0 | 0 | 0 | 0 | 0 | 0 | 0 | 0 | 0 | 0 | 0 | 0 | 0 | 0 | 0 | 0 | 0 | 0 | 0 | 0 | 0 | 0 | 1 | 1 | 1 | 1 | 0 | 0 | 1 | 1 | 1 | 1 | 1 | 1 |
| Aves | *Ciconia ciconia* | 0 | 0 | 1 | 1 | 1 | 1 | 1 | 0 | 0 | 1 | 1 | 1 | 0 | 1 | 1 | 1 | 0 | 0 | 0 | 1 | 1 | 1 | 1 | 1 | 1 | 1 | 1 | 1 | 0 | 0 | 0 | 0 | 0 | 0 | 0 | 0 | 0 | 0 | 0 | 0 |
| | *Colius striatus* | 0 | 0 | 0 | 0 | 0 | 0 | 0 | 0 | 0 | 0 | 0 | 0 | 0 | 0 | 0 | 0 | 0 | 0 | 0 | 0 | 0 | 0 | 0 | 0 | 0 | 0 | 0 | 0 | 1 | 1 | 1 | 1 | 1 | 1 | 1 | 1 | 1 | 1 | 1 | 1 |
| | *Goura sclaterii* | 0 | 0 | 0 | 0 | 0 | 0 | 0 | 0 | 0 | 0 | 0 | 0 | 0 | 0 | 0 | 0 | 0 | 0 | 0 | 0 | 0 | 0 | 0 | 0 | 0 | 0 | 0 | 0 | 1 | 1 | 1 | 1 | 1 | 1 | 1 | 1 | 1 | 1 | 1 | 1 |
| | *Numida meleagris* | 1 | 0 | 1 | 1 | 1 | 1 | 1 | 1 | 1 | 1 | 1 | 1 | 0 | 0 | 0 | 1 | 0 | 0 | 1 | 0 | 0 | 1 | 0 | 1 | 1 | 1 | 1 | 1 | 0 | 0 | 0 | 0 | 0 | 0 | 0 | 0 | 0 | 0 | 0 | 0 |
| | *Gallus gallus* | 1 | 1 | 1 | 1 | 1 | 1 | 0 | 0 | 1 | 1 | 1 | 1 | 1 | 1 | 1 | 1 | 1 | 1 | 1 | 1 | 1 | 1 | 1 | 1 | 1 | 1 | 1 | 1 | 1 | 1 | 1 | 1 | 0 | 0 | 0 | 0 | 0 | 0 | 0 | 0 |
| | *Zapornia flavirostra* | 0 | 0 | 0 | 0 | 0 | 0 | 0 | 0 | 0 | 0 | 0 | 0 | 0 | 0 | 0 | 0 | 0 | 0 | 0 | 0 | 0 | 0 | 0 | 0 | 0 | 0 | 0 | 0 | 0 | 0 | 0 | 0 | 0 | 0 | 0 | 1 | 0 | 0 | 0 | 0 |
| | *Lonchura oryzivora* | 0 | 0 | 0 | 0 | 0 | 0 | 0 | 0 | 0 | 0 | 0 | 0 | 0 | 0 | 0 | 0 | 0 | 0 | 0 | 0 | 0 | 0 | 0 | 0 | 0 | 0 | 0 | 0 | 0 | 0 | 0 | 0 | 1 | 1 | 1 | 1 | 1 | 0 | 1 | 1 |
| | *Ploceus nigricolis* | 0 | 0 | 0 | 0 | 0 | 0 | 0 | 0 | 0 | 0 | 0 | 0 | 0 | 0 | 0 | 0 | 0 | 0 | 0 | 0 | 0 | 0 | 0 | 0 | 0 | 0 | 0 | 0 | 0 | 0 | 0 | 0 | 0 | 1 | 1 | 1 | 1 | 1 | 1 | 1 |
| | *Ramphastos toco* | 0 | 0 | 0 | 0 | 0 | 0 | 0 | 0 | 0 | 0 | 0 | 0 | 0 | 0 | 0 | 0 | 0 | 0 | 0 | 0 | 0 | 0 | 0 | 0 | 0 | 0 | 0 | 0 | 1 | 0 | 0 | 0 | 0 | 0 | 1 | 1 | 0 | 0 | 0 | 0 |
| | *Nestor notabilis* | 0 | 0 | 0 | 0 | 0 | 0 | 0 | 0 | 0 | 0 | 0 | 0 | 0 | 0 | 0 | 0 | 0 | 0 | 0 | 0 | 1 | 1 | 0 | 0 | 1 | 1 | 1 | 1 | 0 | 0 | 0 | 0 | 0 | 0 | 0 | 0 | 0 | 0 | 0 | 0 |
| | *Psittacus erithacus* | 1 | 0 | 1 | 1 | 0 | 0 | 0 | 0 | 1 | 0 | 0 | 0 | 0 | 0 | 0 | 0 | 0 | 0 | 0 | 0 | 0 | 0 | 0 | 0 | 0 | 0 | 0 | 0 | 0 | 0 | 0 | 0 | 0 | 0 | 0 | 0 | 0 | 0 | 0 | 0 |
| | *Struthio camelus* | 0 | 0 | 1 | 0 | 0 | 0 | 0 | 0 | 1 | 1 | 1 | 1 | 0 | 0 | 1 | 1 | 1 | 1 | 1 | 1 | 1 | 1 | 1 | 1 | 1 | 1 | 1 | 1 | 0 | 0 | 0 | 0 | 0 | 0 | 0 | 0 | 0 | 0 | 0 | 0 |
| | *Upupa epops* | 0 | 0 | 0 | 0 | 0 | 0 | 0 | 0 | 0 | 0 | 0 | 0 | 0 | 0 | 0 | 0 | 0 | 0 | 0 | 0 | 0 | 0 | 0 | 0 | 0 | 0 | 0 | 0 | 1 | 1 | 1 | 1 | 0 | 0 | 0 | 0 | 0 | 0 | 0 | 0 |
| Mammalia | *Aepyceros melampus* | 1 | 0 | 0 | 1 | 1 | 1 | 0 | 0 | 1 | 0 | 0 | 0 | 0 | 0 | 1 | 1 | 1 | 0 | 1 | 1 | 0 | 1 | 1 | 0 | 1 | 1 | 0 | 1 | 0 | 0 | 0 | 0 | 0 | 0 | 0 | 0 | 0 | 0 | 0 | 0 |
| | *Bos taurus* | 0 | 1 | 1 | 1 | 1 | 0 | 0 | 0 | 0 | 0 | 0 | 0 | 1 | 1 | 0 | 0 | 0 | 0 | 1 | 1 | 1 | 1 | 1 | 0 | 1 | 1 | 0 | 1 | 1 | 1 | 1 | 1 | 0 | 0 | 0 | 1 | 1 | 1 | 1 | 1 |
| | *Capra hircus* | 0 | 0 | 0 | 0 | 0 | 0 | 0 | 0 | 0 | 0 | 0 | 0 | 0 | 0 | 0 | 0 | 0 | 0 | 0 | 1 | 0 | 1 | 0 | 0 | 1 | 0 | 0 | 0 | 1 | 0 | 0 | 1 | 0 | 0 | 0 | 0 | 0 | 1 | 0 | 0 |
| | *Cephalophus natalensis* | 1 | 1 | 1 | 1 | 1 | 1 | 1 | 1 | 1 | 1 | 1 | 1 | 0 | 0 | 0 | 0 | 0 | 0 | 0 | 0 | 0 | 0 | 0 | 0 | 0 | 0 | 1 | 0 | 0 | 0 | 0 | 0 | 0 | 0 | 0 | 0 | 0 | 0 | 0 | 0 |

| | | | | | | | | | | | | | | | | | | | | | | | | | | | | | | | | | | | | | | | | | |
|---|---|---|---|---|---|---|---|---|---|---|---|---|---|---|---|---|---|---|---|---|---|---|---|---|---|---|---|---|---|---|---|---|---|---|---|---|---|---|---|---|---|
| | *Damaliscus pygargus* | 0 | 0 | 0 | 0 | 0 | 0 | 0 | 0 | 0 | 0 | 0 | 0 | 0 | 0 | 1 | 1 | 0 | 0 | 0 | 1 | 0 | 1 | 0 | 0 | 1 | 1 | 0 | 0 | 0 | 0 | 0 | 0 | 0 | 0 | 0 | 0 | 0 | 0 | 0 | 0 |
| | *Hippotragus niger* | 1 | 0 | 1 | 1 | 0 | 1 | 0 | 0 | 1 | 0 | 1 | 1 | 1 | 1 | 1 | 1 | 1 | 1 | 1 | 1 | 0 | 0 | 0 | 0 | 1 | 1 | 1 | 1 | 0 | 0 | 0 | 0 | 0 | 0 | 0 | 0 | 0 | 0 | 0 | 0 |
| | *Giraffa camelopardalis* | 1 | 0 | 1 | 1 | 1 | 1 | 1 | 0 | 1 | 1 | 1 | 1 | 0 | 1 | 1 | 1 | 0 | 1 | 1 | 1 | 1 | 1 | 1 | 1 | 1 | 1 | 1 | 1 | 0 | 0 | 0 | 0 | 0 | 0 | 0 | 0 | 0 | 0 | 0 | 0 |
| | *Okapia johnstoni* | 1 | 1 | 1 | 1 | 1 | 1 | 1 | 1 | 1 | 1 | 1 | 1 | 1 | 1 | 1 | 1 | 1 | 0 | 1 | 1 | 0 | 0 | 0 | 0 | 0 | 0 | 1 | 1 | 1 | 0 | 1 | 0 | 0 | 1 | 0 | 0 | 0 | 0 | 0 | 0 |
| | *Sus scrofa* | 1 | 1 | 1 | 1 | 1 | 1 | 1 | 1 | 1 | 1 | 1 | 1 | 1 | 1 | 1 | 1 | 1 | 1 | 1 | 1 | 1 | 1 | 1 | 1 | 1 | 1 | 1 | 1 | 1 | 1 | 1 | 1 | 1 | 0 | 1 | 1 | 1 | 1 | 1 | 1 |
| | *Canis lupus* | 1 | 1 | 1 | 1 | 1 | 1 | 1 | 1 | 0 | 0 | 0 | 0 | 1 | 1 | 1 | 1 | 1 | 1 | 1 | 1 | 1 | 1 | 1 | 1 | 1 | 1 | 1 | 1 | 1 | 1 | 1 | 1 | 0 | 1 | 1 | 1 | 1 | 1 | 1 | 1 |
| | *Felis catus* | 0 | 0 | 0 | 0 | 0 | 0 | 0 | 0 | 0 | 0 | 0 | 0 | 0 | 1 | 0 | 0 | 0 | 0 | 1 | 0 | 0 | 0 | 0 | 0 | 0 | 0 | 0 | 0 | 0 | 0 | 0 | 0 | 0 | 0 | 0 | 0 | 0 | 1 | 0 | 0 |
| | *Mungus mungo* | 0 | 0 | 0 | 0 | 0 | 0 | 0 | 0 | 0 | 0 | 0 | 0 | 0 | 0 | 1 | 1 | 1 | 0 | 1 | 1 | 1 | 1 | 0 | 0 | 1 | 1 | 1 | 1 | 0 | 0 | 0 | 0 | 0 | 0 | 0 | 0 | 0 | 0 | 0 | 0 |
| | *Tolypeutes matacus* | 0 | 0 | 0 | 0 | 0 | 0 | 0 | 0 | 0 | 0 | 0 | 0 | 0 | 0 | 0 | 0 | 0 | 0 | 0 | 0 | 0 | 0 | 0 | 0 | 0 | 0 | 0 | 0 | 0 | 0 | 0 | 1 | 0 | 0 | 0 | 0 | 0 | 0 | 0 | 0 |
| | *Macropus giganteus* | 0 | 0 | 0 | 0 | 0 | 1 | 0 | 0 | 0 | 1 | 0 | 1 | 0 | 0 | 1 | 0 | 0 | 0 | 0 | 0 | 0 | 0 | 0 | 0 | 0 | 0 | 0 | 0 | 1 | 0 | 0 | 1 | 0 | 0 | 0 | 0 | 0 | 0 | 0 | 0 |
| | *Lepus europaeus* | 0 | 0 | 0 | 0 | 0 | 0 | 0 | 0 | 0 | 0 | 0 | 0 | 0 | 0 | 0 | 0 | 0 | 0 | 0 | 0 | 0 | 0 | 0 | 0 | 1 | 0 | 0 | 0 | 0 | 0 | 0 | 0 | 0 | 0 | 0 | 0 | 0 | 0 | 0 | 0 |
| | *Oryctolagus cuniculus* | 0 | 0 | 0 | 0 | 0 | 0 | 0 | 0 | 0 | 0 | 0 | 0 | 0 | 0 | 1 | 1 | 0 | 0 | 1 | 1 | 0 | 1 | 0 | 0 | 1 | 1 | 0 | 0 | 0 | 0 | 0 | 0 | 0 | 0 | 0 | 0 | 0 | 1 | 0 | 0 |
| | *Equus caballus* | 0 | 0 | 0 | 1 | 0 | 0 | 0 | 0 | 0 | 0 | 0 | 0 | 0 | 0 | 0 | 1 | 1 | 0 | 1 | 1 | 1 | 0 | 1 | 0 | 1 | 1 | 1 | 1 | 0 | 1 | 0 | 1 | 0 | 0 | 1 | 1 | 0 | 0 | 0 | 0 |
| | *Equus quagga* | 1 | 0 | 0 | 0 | 0 | 0 | 0 | 0 | 0 | 0 | 0 | 0 | 0 | 0 | 1 | 1 | 1 | 1 | 0 | 1 | 0 | 0 | 0 | 0 | 1 | 1 | 1 | 0 | 0 | 0 | 0 | 0 | 0 | 0 | 0 | 0 | 0 | 0 | 0 | 0 |
| | *Ceratotherium simum* | 1 | 1 | 1 | 1 | 0 | 0 | 0 | 0 | 1 | 1 | 1 | 1 | 0 | 1 | 1 | 1 | 0 | 0 | 1 | 1 | 1 | 1 | 1 | 1 | 1 | 1 | 1 | 1 | 0 | 0 | 0 | 0 | 0 | 0 | 0 | 0 | 0 | 0 | 0 | 0 |
| | *Choloepus didactylus* | 0 | 0 | 0 | 0 | 0 | 0 | 0 | 0 | 0 | 0 | 0 | 0 | 0 | 0 | 0 | 0 | 0 | 0 | 0 | 0 | 0 | 0 | 0 | 0 | 0 | 0 | 0 | 0 | 1 | 1 | 0 | 1 | 1 | 1 | 1 | 1 | 1 | 1 | 1 | 1 |
| | *Leontopithecus rosalia* | 0 | 0 | 0 | 0 | 0 | 0 | 0 | 0 | 0 | 0 | 0 | 0 | 0 | 0 | 0 | 0 | 0 | 0 | 0 | 0 | 0 | 0 | 0 | 0 | 0 | 0 | 0 | 0 | 0 | 1 | 0 | 0 | 0 | 0 | 0 | 0 | 0 | 0 | 0 | 0 |
| | *Lemur catta* | 0 | 0 | 0 | 0 | 0 | 0 | 0 | 0 | 0 | 0 | 0 | 0 | 0 | 0 | 0 | 0 | 0 | 0 | 0 | 0 | 0 | 1 | 0 | 0 | 0 | 0 | 0 | 0 | 0 | 0 | 0 | 0 | 0 | 0 | 0 | 0 | 0 | 0 | 0 | 0 |
| | *Elephas maximus* | 0 | 0 | 0 | 0 | 0 | 0 | 0 | 0 | 0 | 0 | 0 | 0 | 0 | 0 | 0 | 0 | 0 | 0 | 0 | 0 | 0 | 0 | 0 | 0 | 0 | 0 | 0 | 0 | 0 | 1 | 0 | 0 | 0 | 0 | 1 | 0 | 0 | 0 | 1 | 0 |
| | *Cavia porcellus* | 0 | 0 | 0 | 0 | 0 | 0 | 0 | 0 | 0 | 0 | 0 | 0 | 0 | 0 | 0 | 0 | 0 | 0 | 0 | 0 | 0 | 1 | 0 | 0 | 0 | 0 | 0 | 0 | 0 | 0 | 0 | 0 | 0 | 0 | 0 | 0 | 0 | 0 | 0 | 0 |
| | *Arvicola amphibius* | 0 | 0 | 0 | 1 | 0 | 0 | 0 | 0 | 0 | 0 | 0 | 0 | 0 | 0 | 0 | 0 | 0 | 0 | 0 | 0 | 0 | 0 | 0 | 0 | 0 | 0 | 0 | 0 | 0 | 0 | 0 | 0 | 0 | 0 | 0 | 0 | 0 | 0 | 0 | 0 |
| | *Apodemus flavicollis* | 0 | 0 | 0 | 0 | 0 | 0 | 0 | 0 | 0 | 0 | 0 | 0 | 1 | 0 | 0 | 0 | 0 | 0 | 1 | 0 | 0 | 0 | 0 | 0 | 0 | 0 | 0 | 0 | 0 | 0 | 0 | 1 | 1 | 0 | 1 | 0 | 1 | 1 | 1 | 0 |
| | *Mus musculus* | 1 | 0 | 1 | 1 | 1 | 1 | 0 | 1 | 1 | 1 | 1 | 1 | 0 | 0 | 1 | 1 | 1 | 0 | 1 | 1 | 1 | 1 | 1 | 0 | 1 | 1 | 0 | 1 | 0 | 0 | 0 | 0 | 0 | 0 | 1 | 1 | 1 | 1 | 1 | 1 |
| | *Rattus norvegicus* | 0 | 0 | 0 | 1 | 0 | 0 | 0 | 0 | 0 | 0 | 0 | 0 | 0 | 0 | 0 | 0 | 0 | 0 | 1 | 1 | 0 | 0 | 0 | 0 | 1 | 1 | 0 | 1 | 0 | 0 | 0 | 0 | 0 | 0 | 0 | 0 | 0 | 0 | 0 | 0 |
| | *Sciurus vulgaris* | 0 | 0 | 0 | 0 | 0 | 0 | 0 | 0 | 0 | 0 | 0 | 0 | 1 | 0 | 0 | 1 | 0 | 1 | 0 | 0 | 0 | 0 | 0 | 0 | 0 | 0 | 0 | 0 | 0 | 0 | 0 | 0 | 0 | 0 | 0 | 0 | 0 | 0 | 0 | 0 |
| | *Tragulus javanicus* | 0 | 0 | 0 | 0 | 0 | 0 | 0 | 0 | 0 | 0 | 0 | 0 | 0 | 0 | 0 | 0 | 0 | 0 | 0 | 0 | 0 | 0 | 0 | 0 | 0 | 0 | 0 | 0 | 0 | 1 | 0 | 1 | 0 | 0 | 0 | 0 | 0 | 0 | 0 | 0 |
| Reptilia | *Acrantophis dumerili* | 0 | 0 | 0 | 0 | 0 | 0 | 0 | 0 | 0 | 0 | 0 | 0 | 0 | 0 | 0 | 0 | 0 | 0 | 0 | 0 | 0 | 0 | 0 | 0 | 0 | 0 | 0 | 0 | 0 | 0 | 0 | 0 | 0 | 0 | 0 | 1 | 0 | 0 | 0 | 0 |
| Total detections per replicate | | 13 | 7 | 15 | 17 | 11 | 12 | 8 | 6 | 13 | 11 | 11 | 12 | 8 | 10 | 17 | 18 | 11 | 8 | 18 | 20 | 12 | 18 | 11 | 8 | 21 | 20 | 15 | 17 | 15 | 14 | 10 | 17 | 7 | 8 | 13 | 15 | 12 | 13 | 12 | 10 |
| Total unique detections | | 7 | 1 | 2 | 3 | 1 | 2 | 3 | 1 | 4 | 2 | | 1 | 2 | 4 | 2 | 3 | 4 | 2 | 3 | 4 | 1 | 7 | 4 | 1 | 2 | 1 | 2 | 4 | 5 | 4 | 1 | 8 | 2 | 3 | 2 | 4 | 1 | 3 | 1 | |
| Total shared detections | | 6 | | 13 | | 10 | | 5 | | 9 | | 11 | | 6 | | 15 | | 6 | | 15 | | 11 | | 7 | | 19 | | 13 | | 10 | | 9 | | 5 | | 11 | | 10 | | 10 | |

**Supplementary Table 3.** List of non-human vertebrates species detected in air samples (n=14) sampling in open air, and by using the 16S mammal (m) and the 12S. Detections range from non-detected (0) to detection in both replicates (2) in the different samplers used: using water vacuum (WV) for 30 and 60 min, particle filter sampler with the 24V fan (MF) ran during 30, 60 and 5hrs, and particle filter sampler with the 5V fan(SF) ran during 30 hrs. Samples collected outside among the outdoor animal enclosures. Information regarding the presence of the species in the southern area of the zoo, and in the rest of the zoo, is provided.

| | | | | | | September | | December | | | | | | | |
|---|---|---|---|---|---|---|---|---|---|---|---|---|---|---|---|
| Class | Order | Family | *Species* | Common name | Primer | WV_30 min | WV_60 min | WV_30 min | WV_60 min | MF_30 min | MF_60 min | MF_5hrs | SF_30rhs | Present in southern area | Zoo animal |
| Actinopterygii | Clupeiformes | Clupeidae | NA | NA | v | 0 | 0 | 0 | 0 | 0 | 0 | 1 | 0 | NA | NA |
| | Osmeriformes | Osmeridae | *Osmerus eperlanus* | European smelt | v | 0 | 1 | 0 | 1 | 0 | 0 | 1 | 0 | No | Food |
| Aves | Anseriformes | Anatidae | NA | NA | v | 0 | 2 | 1 | 1 | 0 | 0 | 2 | 0 | NA | NA |
| | Ciconiiformes | Ciconiidae | *Ciconia ciconia* | White stork | v | 1 | 2 | 0 | 1 | 2 | 2 | 2 | 2 | Yes | Yes |
| | Columbiformes | Columbidae | NA | NA | v | 2 | 2 | 2 | 2 | 2 | 2 | 2 | 0 | NA | NA |
| | Galliformes | Numididae | *Numida meleagris* | Helmeted guineafowl | v | 0 | 1 | 0 | 1 | 1 | 1 | 2 | 2 | Yes | Yes |
| | | Phasianidae | *Gallus gallus* | Domestic fowl | v | 2 | 2 | 2 | 2 | 2 | 2 | 2 | 2 | Yes | Yes |
| | Passeriformes | Corvidae | *Corvus* sp. | Corvid | m | 0 | 0 | 0 | 1 | 0 | 0 | 0 | 0 | No | No |
| | | Corvidae | NA | NA | v | 0 | 2 | 0 | 0 | 0 | 1 | 0 | 0 | NA | NA |
| | | NA | NA | NA | v | 0 | 2 | 1 | 1 | 1 | 0 | 2 | 0 | NA | NA |
| | Psittaciformes | Psittacidae | *Nestor notabilis* | Kea | v | 0 | 0 | 0 | 0 | 2 | 0 | 2 | 2 | Yes | Yes |
| | Struthioniformes | Struthionidae | *Struthio camelus* | Common ostrich | v | 0 | 2 | 2 | 2 | 2 | 2 | 2 | 2 | Yes | Yes |
| Mammalia | Artiodactyla | Bovidae | *Aepyceros melampus* | Impala | m/v | 0 | 2 | 1 | 2 | 1 | 1 | 2 | 1 | Yes | Yes |
| | | | *Bos taurus* | Domestic cow/ox | m/v | 2 | 0 | 0 | 2 | 2 | 1 | 2 | 1 | Yes | Yes |
| | | | *Capra hircus* | Pygmy goat | m | 0 | 0 | 0 | 1 | 1 | 0 | 1 | 0 | Yes | Yes |
| | | | *Cephalophus natalensis* | Red forest duiker | m | 0 | 0 | 0 | 0 | 0 | 0 | 0 | 1 | Yes | Yes |
| | | | *Damaliscus pygargus* | Bontebok / blesbok | m | 0 | 2 | 0 | 1 | 1 | 0 | 2 | 0 | Yes | Yes |
| | | | *Hippotragus niger* | Sable antelope | m/v | 2 | 2 | 2 | 2 | 0 | 0 | 2 | 2 | Yes | Yes |
| | | Giraffidae | *Giraffa camelopardalis* | Giraffe | m/v | 1 | 2 | 1 | 2 | 1 | 2 | 2 | 2 | Yes | Yes |
| | | | *Okapia johnstoni* | Okapi | m/v | 2 | 2 | 2 | 2 | 0 | 1 | 1 | 2 | Yes | Yes |

| | | | | | | | | | | | | | | |
|---|---|---|---|---|---|---|---|---|---|---|---|---|---|---|
| | | Suidae | *Sus scrofa* | Domestic pig | m/v | 2 | 2 | 2 | 2 | 2 | 2 | 2 | 2 | Yes | Yes |
| | Carnivora | Canidae | *Canis lupus* | Dog / gray wolf | m/v | 2 | 2 | 2 | 2 | 2 | 2 | 2 | 2 | No | No |
| | | Felidae | *Felis catus* | Domestic cat | m | 1 | 0 | 0 | 1 | 0 | 0 | 0 | 0 | No | No |
| | | Herpestidae | *Mungos mungo* | Banded mongoose | m/v | 0 | 2 | 1 | 2 | 2 | 0 | 2 | 2 | Yes | Yes |
| | Diprotodontia | Macropodidae | *Macropus giganteus* | Eastern grey kangaroo | v | 0 | 1 | 0 | 0 | 0 | 0 | 0 | 0 | No | Yes |
| | Lagomorpha | Leporidae | *Lepus europaeus* | European hare | m | 0 | 0 | 0 | 0 | 0 | 0 | 1 | 0 | No | No |
| | | Leporidae | *Oryctolagus cuniculus* | Domestic rabbit | m | 0 | 2 | 0 | 2 | 1 | 0 | 2 | 0 | Yes | Yes |
| | Perissodactyla | Equidae | *Equus caballus* | Horse | m | 0 | 1 | 1 | 2 | 1 | 1 | 2 | 2 | Yes | Yes |
| | | | *Equus quagga* | Plains zebra | m/v | 0 | 2 | 2 | 1 | 0 | 0 | 2 | 1 | Yes | Yes |
| | | Rhinocerotidae | *Ceratotherium simum* | White rhinoceros | m/v | 1 | 2 | 0 | 2 | 2 | 2 | 2 | 2 | Yes | Yes |
| | Primates | Lemuridae | *Lemur catta* | Ring-tailed lemur | m | 0 | 0 | 0 | 0 | 1 | 0 | 0 | 0 | Yes | Yes |
| | Rodentia | Caviidae | *Cavia porcellus* | Domestic guinea pig | m | 0 | 0 | 0 | 0 | 1 | 0 | 0 | 0 | Yes | Yes |
| | | Cricetidae | NA | NA | m | 0 | 1 | 0 | 0 | 0 | 0 | 0 | 0 | NA | NA |
| | | Muridae | *Apodemus flavicollis* | Yellow-necked mouse | m | 1 | 0 | 0 | 1 | 0 | 0 | 0 | 0 | No | No |
| | | | *Apodemus* sp. | NA | v | 1 | 1 | 0 | 0 | 0 | 0 | 0 | 0 | NA | NA |
| | | | *Mus musculus* | House mouse | m/v | 0 | 2 | 1 | 2 | 2 | 1 | 2 | 1 | No | Yes |
| | | | *Rattus norvegicus* | Brown rat | m | 0 | 0 | 0 | 2 | 0 | 0 | 2 | 1 | No | Yes |
| | | Sciuridae | *Sciurus vulgaris* | Eurasian red squirrel | m | 1 | 1 | 1 | 0 | 0 | 0 | 0 | 0 | No | No |

**Supplementary Table 4.** List of non-human vertebrate species detected in air samples (n=12) sampling in the Rainforest House within the Tropical House, and by using the 16S mammal (m) and the 12S vertebrate (v) metabarcoding primers. Detections range from non-detected (0) to detection in both replicates (2) in the different samplers used: using water vacuum (WV) for 30 and 60 min, particle filter sampler with the 24V fan (MF) ran during 30, 60 and 5hrs, and particle filter sampler with the 5V fan(SF) ran during 30 hrs. Samples collected outside among the outdoor animal enclosures. Information regarding the presence of the species in the Tropical House, and in the rest of the zoo, is provided.

| Class | Order | Family | Species | Common name | Primer | WV_30min | WV_60min | MF_30min | MF_60min | MF_5hrs | SF_30rhs | Present in Tropical House | Zoo animal |
|---|---|---|---|---|---|---|---|---|---|---|---|---|---|
| Actinopterygii | Cypriniformes | Cyprinidae | *Rutilus rutilus* | Common roach | v | 1 | 0 | 0 | 0 | 0 | 0 | No | Food |
| | Cyprinodontiformes | Poeciliidae | *Poecilia reticulata* | Guppy | v | 2 | 2 | 2 | 1 | 1 | 0 | Yes | Yes |
| | Salmoniformes | Salmonidae | *Salmo salar* | Atlantic salmon | v | 0 | 1 | 0 | 0 | 0 | 0 | No | Food |
| Amphibia | Anura | Bufonidae | *Duttaphrynus melanostictus* | Asian common toad | m | 2 | 2 | 0 | 2 | 2 | 2 | Yes | Yes |
| Aves | Anseriformes | Anatidae | NA | NA | v | 2 | 1 | 1 | 1 | 0 | 0 | NA | NA |
| | Coliiformes | Coliidae | *Colius striatus* | Speckled mousebird | m/v | 2 | 2 | 2 | 2 | 2 | 2 | Yes | Yes |
| | Columbiformes | Columbidae | *Goura sclaterii* | Sclater's crowned-pigeon | m/v | 2 | 2 | 2 | 2 | 2 | 2 | Yes | Yes |
| | | Columbidae | NA | NA | v | 2 | 2 | 0 | 0 | 0 | 0 | Yes | Yes |
| | Galliformes | Phasianidae | *Gallus gallus* | Domestic fowl | v | 2 | 2 | 0 | 0 | 0 | 0 | Food | Food |
| | Gruiformes | Rallidae | *Zapornia flavirostra* | Black crake | m | 0 | 0 | 0 | 1 | 0 | 0 | Yes | Yes |
| | Passeriformes | Estrildidae | *Lonchura oryzivora* | Javan sparrow | m | 0 | 0 | 2 | 2 | 1 | 2 | Yes | Yes |
| | | Ploceidae | *Ploceus sp.* | NA | v | 1 | 1 | 2 | 2 | 1 | 1 | Yes | Yes |
| | | | *Ploceus nigricollis* | Black-necked weaver | m | 0 | 0 | 1 | 2 | 2 | 2 | Yes | Yes |
| | | Sturnidae | *Sturnus sp.* | Starling | m | 0 | 0 | 0 | 1 | 1 | 1 | No | No |
| | | Corvidae | NA | NA | v | 1 | 2 | 0 | 0 | 0 | 0 | NA | NA |
| | | NA | NA | NA | v | 2 | 2 | 2 | 2 | 2 | 2 | Yes | NA |
| | Piciformes | Ramphastidae | *Ramphastos toco* | Toco toucan | v | 1 | 0 | 0 | 2 | 0 | 0 | Yes | Yes |
| | Upupiformes | Upupidae | *Upupa epops* | Eurasian hoopoe | v | 2 | 2 | 0 | 0 | 0 | 0 | Yes | Yes |
| Mammalia | Artiodactyla | Bovidae | *Bos taurus* | Domestic cow/ox | m/v | 2 | 2 | 0 | 1 | 2 | 2 | No | Yes |
| | | | *Capra hircus* | Pygmy goat | m | 1 | 1 | 0 | 0 | 1 | 0 | No | Yes |
| | | Giraffidae | *Okapia johnstoni* | Okapi | m/v | 2 | 2 | 1 | 1 | 2 | 0 | No | Yes |
| | | Suidae | *Sus scrofa* | Domestic pig | m/v | 2 | 2 | 1 | 2 | 2 | 2 | No | Yes |

| | | | | | | | | | | | | | |
|---|---|---|---|---|---|---|---|---|---|---|---|---|---|
| | | Tragulidae | *Tragulus javanicus* | Java mouse-deer | m | 1 | 1 | 0 | 0 | 0 | 0 | Yes | Yes |
| | Carnivora | Canidae | *Canis lupus* | Dog / gray wolf | m/v | 2 | 2 | 1 | 2 | 2 | 2 | No | No |
| | | Felidae | *Felis catus* | Domestic cat | m | 0 | 0 | 0 | 0 | 1 | 0 | No | No |
| | Cingulata | Chlamyphoridae | *Tolypeutes matacus* | Southern three-banded armadillo | m | 0 | 1 | 0 | 0 | 0 | 0 | Yes | Yes |
| | Diprotodontia | Macropodidae | *Macropus giganteus* | Eastern grey kangaroo | m/v | 1 | 1 | 0 | 0 | 0 | 0 | No | Yes |
| | Lagomorpha | Leporidae | *Oryctolagus cuniculus* | Domestic rabbit | m | 0 | 0 | 0 | 0 | 1 | 0 | Food | Yes |
| | Perissodactyla | Equidae | *Equus caballus* | Horse | m | 1 | 1 | 0 | 2 | 0 | 0 | No | Yes |
| | Pilosa | Megalonychidae | *Choloepus didactylus* | Linne's two-toed sloth | m | 2 | 1 | 2 | 2 | 2 | 2 | Yes | Yes |
| | Primates | Cebidae | *Leontopithecus rosalia* | Golden lion tamarin | m | 1 | 0 | 0 | 0 | 0 | 0 | Yes | Yes |
| | Proboscidea | Elephantidae | *Elephas maximus* | Asian elephant | m | 1 | 0 | 0 | 1 | 0 | 1 | No | Yes |
| | Rodentia | Muridae | *Apodemus flavicollis* | Yellow-necked mouse | m | 0 | 1 | 1 | 1 | 2 | 1 | No | No |
| | | | *Apodemus sp.* | NA | v | 0 | 0 | 1 | 0 | 0 | 0 | No | No |
| | | Muridae | *Mus musculus* | House mouse | m | 0 | 0 | 0 | 2 | 2 | 2 | No | Yes |
| | | NA | NA | NA | m | 0 | 1 | 0 | 0 | 0 | 0 | NA | NA |
| Reptilia | Squamata | Boidae | *Acrantophis dumerili* | Dumeril's ground boa | m | 0 | 0 | 0 | 1 | 0 | 0 | Yes | Yes |

**Supplementary Table 5.** List of terrestrial vertebrate species present in the Tropical House in Copenhagen Zoo, specifying their presence inside the Rainforest House where air samples were collected.

| Class | Species | Common name | Present in Rainforest House |
|---|---|---|---|
| Amphibia | *Bombina variegata* | European yellow-bellied toad | No |
| | *Dendrobates auratus* | Green and black poison frog | No |
| | *Dendrobates leucomelas* | Yellow-headed poison frog | No |
| | *Dendrobates tinctorius* | Dyeing poison frog | No |
| | *Duttaphrynus melanostictus* | Asian common toad | Yes |
| | *Dyscophus guineti* | Sambava tomato frog | Yes |
| | *Phyllobates terribilis* | Golden poison dart frog | No |
| | *Phyllobates vittatus* | Golfodulcean poison dart frog | No |
| | *Polypedates dennysi* | Denny's tree frog | No |
| | *Theloderma corticale* | Tonkin bug-eyed frog | No |
| | *Trachycephalus resinifictrix* | Mission golden-eyed tree frog | No |
| | *Tylototriton shanjing* | Emperor newt | No |
| Aves | *Amazona aestiva* | Blue-fronted amazon | No |
| | *Amazona amazonica* | Orange-winged amazon | No |
| | *Amazona auropalliata* | Yellow-naped amazon | No |
| | *Amazona barbadensis* | Yellow-shouldered amazon | No |
| | *Amazona leucocephala* | Cuban amazon | No |
| | *Amazona ochrocephala* | Yellow-crowned amazon | No |
| | *Ara ararauna* | Blue-and-yellow macaw | No |
| | *Bycanistes bucinator* | Trumpeter hornbill | No |
| | *Charmosyna stellae* | Stella's lorikeet | No |
| | *Cinnyricinclus leucogaster* | Violet-backed starling | Yes |
| | *Colius striatus* | Speckled mousebird | No |
| | *Colius striatus* | Speckled mousebird | Yes |
| | *Cotinga cayana* | Spangled cotinga | Yes |
| | *Goura sclaterii* | Sclater's crowned-pigeon | Yes |

| | | | |
|---|---|---|---|
| | *Irena puella* | Fairy bluebird | Yes |
| | *Kittacincla malabarica* | White-rumped shama | Yes |
| | *Leiothrix lutea* | Red-billed leiothrix | Yes |
| | *Leucopsar rothschildi* | Bali myna | No |
| | *Lonchura oryzivora* | Javan sparrow | No |
| | *Loriculus galgulus* | Blue-crowned parrot | No |
| | *Musophaga violacea* | Violet turaco | Yes |
| | *Nettapus auritus* | African pygmy goose | Yes |
| | *Otidiphaps aruensis* | White-naped pheasant-pigeon | Yes |
| | *Paroaria dominicana* | Red-cowled cardinal | Yes |
| | *Ploceus nigricollis* | Black-necked weaver | Yes |
| | *Pycnonotus jocosus* | Red-whiskered bulbul | Yes |
| | *Ramphastos toco* | Toco toucan | No |
| | *Ramphocelus bresilius* | Brazilian tanager | No |
| | *Tiaris canora* | Cuban grassquit | No |
| | *Trachyphonus erythrocephalus* | Red-and-yellow barbet | Yes |
| | *Upupa epops* | Eurasian hoopoe | Yes |
| | *Volatinia jacarina* | Blue-black grassquit | No |
| | *Zapornia flavirostra* | Black crake | Yes |
| Mammal | *Choloepus didactylus* | Linne's two-toed sloth | Yes |
| | *Leontopithecus rosalia* | Golden lion tamarin | No |
| | *Tolypeutes matacus* | Southern three-banded armadillo | Yes |
| | *Tragulus javanicus* | Java mouse-deer | No |
| Reptilia | *Acrantophis dumerili* | Dumeril's ground boa | Yes |
| | *Brachylophus fasciatus* | Lau banded iguana | No |
| | *Carettochelys insculpta* | Fly River turtle | Yes |
| | *Chelonoidis carbonarius* | Red-footed tortoise | Yes |
| | *Chilabothrus subflavus* | Jamaican boa | No |
| | *Crocodylus mindorensis* | Philippine crocodile | No |
| | *Cuora amboinensis* | Southeast Asian box turtle | No |
| | *Cuora flavomarginata* | Yellow-margined box turtle | No |
| | *Dracaena guianensis* | Caiman lizard | No |

| | | |
|---|---|---|
| *Elaphe schrencki* | Amur ratsnake | No |
| *Eublepharis macularius* | Leopard gecko | No |
| *Eunectes murinus* | Green anaconda | No |
| *Lampropeltis getula* | Common kingsnake | No |
| *Lampropeltis triangulum* | Milksnake | No |
| *Lamprophis fuliginosus* | Brown house snake | No |
| *Morelia viridis* | Green tree python | No |
| *Phelsuma grandis* | Giant Madagascar day gecko | No |
| *Python regius* | Royal/ball python | No |
| *Rhacodactylus auriculatus* | New Caledonia bumpy gecko | No |
| *Stenodactylus petrii* | Petrie's gecko | No |
| *Tiliqua gerrardii* | Pink-tongued skink | No |
| *Varanus varius* | Lace monitor | Yes |

**Supplementary Table 6.** List of non-human vertebrate species present in the southern part of the Copenhagen Zoo, used for the logistic model. Detection (1) or non-detection (0) of these species, with the different samplers used: using water vacuum (WV) for 30 and 60 min, particle filter sampler with the 24V fan (MF) ran during 30, 60 and 5hrs, and particle filter sampler with the 5V fan (SF) ran during 30 hrs. Data includes number of individuals per species present in that area, their weight, their total biomass and the distance from the location of that species to the samplers.

| Class | Genus | Species | Detection | WV | MF | SF | WV 30min | WV 60min | MF 30min | MF 60min | MF 5hrs | SF 30hrs | N individuals | Weight (kg) | Biomass | Distance |
|---|---|---|---|---|---|---|---|---|---|---|---|---|---|---|---|---|
| Aves | *Nettapus* | *auritus* | 0 | 0 | 0 | 0 | 0 | 0 | 0 | 0 | 0 | 0 | 4 | 0.275 | 1.1 | 70 |
| Aves | *Bucorvus* | *abyssinicus* | 0 | 0 | 0 | 0 | 0 | 0 | 0 | 0 | 0 | 0 | 2 | 4 | 8 | 70 |
| Aves | *Upupa* | *epops* | 0 | 0 | 0 | 0 | 0 | 0 | 0 | 0 | 0 | 0 | 11 | 0.067 | 0.737 | 70 |
| Aves | *Ciconia* | *ciconia* | 1 | 1 | 1 | 1 | 1 | 1 | 1 | 1 | 1 | 1 | 4 | 3.3 | 13.2 | 22 |
| Aves | *Nesoenas* | *mayeri* | 0 | 0 | 0 | 0 | 0 | 0 | 0 | 0 | 0 | 0 | 1 | 0.36 | 0.36 | 70 |
| Aves | *Numida* | *meleagris* | 1 | 1 | 1 | 1 | 1 | 1 | 1 | 1 | 1 | 1 | 25 | 1.3 | 32.5 | 60 |
| Aves | *Afropavo* | *congensis* | 0 | 0 | 0 | 0 | 0 | 0 | 0 | 0 | 0 | 0 | 4 | 1.35 | 5.4 | 70 |
| Aves | *Tauraco* | *erythrolophus* | 0 | 0 | 0 | 0 | 0 | 0 | 0 | 0 | 0 | 0 | 2 | 0.05 | 0.1 | 70 |
| Aves | *Ploceus* | *capensis* | 0 | 0 | 0 | 0 | 0 | 0 | 0 | 0 | 0 | 0 | 15 | 0.044 | 0.66 | 70 |
| Aves | *Lamprotornis* | *superbus* | 0 | 0 | 0 | 0 | 0 | 0 | 0 | 0 | 0 | 0 | 9 | 0.064 | 0.576 | 70 |
| Aves | *Nestor* | *notabilis* | 1 | 0 | 1 | 1 | 0 | 0 | 1 | 0 | 1 | 1 | 5 | 0.9 | 4.5 | 120 |
| Aves | *Psittacus* | *erithacus* | 0 | 0 | 0 | 0 | 0 | 0 | 0 | 0 | 0 | 0 | 5 | 0.407 | 2.035 | 70 |
| Aves | *Agapornis* | *nigrigenis* | 0 | 0 | 0 | 0 | 0 | 0 | 0 | 0 | 0 | 0 | 95 | 0.04 | 3.8 | 70 |
| Aves | *Struthio* | *camelus* | 1 | 1 | 1 | 1 | 1 | 1 | 1 | 1 | 1 | 1 | 2 | 90 | 180 | 60 |
| Mammalia | *Aepyceros* | *melampus* | 1 | 1 | 1 | 1 | 1 | 1 | 1 | 1 | 1 | 1 | 13 | 50 | 650 | 60 |
| Mammalia | *Bos* | *taurus* | 1 | 1 | 1 | 1 | 1 | 1 | 1 | 1 | 1 | 1 | 4 | 600 | 2400 | 160 |
| Mammalia | *Capra* | *hircus* | 1 | 1 | 1 | 0 | 0 | 1 | 1 | 0 | 1 | 0 | 22 | 20 | 440 | 175 |
| Mammalia | *Cephalophus* | *natalensis* | 1 | 0 | 0 | 1 | 0 | 0 | 0 | 0 | 0 | 1 | 2 | 10 | 20 | 15 |
| Mammalia | *Damaliscus* | *pygargus* | 1 | 1 | 1 | 0 | 0 | 1 | 1 | 0 | 1 | 0 | 3 | 67 | 201 | 40 |
| Mammalia | *Hippotragus* | *niger* | 1 | 1 | 1 | 1 | 1 | 1 | 0 | 0 | 1 | 1 | 10 | 229 | 2290 | 60 |
| Mammalia | *Lama* | *glama* | 0 | 0 | 0 | 0 | 0 | 0 | 0 | 0 | 0 | 0 | 2 | 140 | 280 | 190 |
| Mammalia | *Giraffa* | *camelopardalis* | 1 | 1 | 1 | 1 | 1 | 1 | 1 | 1 | 1 | 1 | 4 | 500 | 2000 | 60 |
| Mammalia | *Okapia* | *johnstoni* | 1 | 1 | 1 | 1 | 1 | 1 | 1 | 1 | 1 | 1 | 2 | 250 | 500 | 40 |
| Mammalia | *Hippopotamus* | *amphibius* | 0 | 0 | 0 | 0 | 0 | 0 | 0 | 0 | 0 | 0 | 5 | 1800 | 9000 | 93 |
| Mammalia | *Sus* | *scrofa* | 1 | 1 | 1 | 1 | 0 | 1 | 1 | 0 | 1 | 0 | 2 | 169 | 338 | 196 |

| | | | | | | | | | | | | | | | | |
|---|---|---|---|---|---|---|---|---|---|---|---|---|---|---|---|---|
| Mammalia | *Caracal* | *caracal* | 0 | 0 | 0 | 0 | 0 | 0 | 0 | 0 | 0 | 0 | 2 | 13 | 26 | 97 |
| Mammalia | *Mungos* | *mungo* | 1 | 1 | 1 | 1 | 1 | 0 | 1 | 0 | 1 | 1 | 6 | 1.25 | 7.5 | 20 |
| Mammalia | *Macropus* | *rufogriseus* | 0 | 0 | 0 | 0 | 0 | 0 | 0 | 0 | 0 | 0 | 12 | 16 | 192 | 93 |
| Mammalia | *Oryctolagus* | *cuniculus* | 1 | 1 | 1 | 0 | 0 | 1 | 1 | 0 | 1 | 1 | 15 | 2 | 30 | 157 |
| Mammalia | *Equus* | *caballus* | 1 | 1 | 1 | 1 | 1 | 1 | 1 | 1 | 1 | 1 | 5 | 250 | 1250 | 160 |
| Mammalia | *Equus* | *quagga* | 1 | 1 | 1 | 1 | 1 | 1 | 0 | 0 | 1 | 1 | 5 | 275 | 1375 | 60 |
| Mammalia | *Ceratotherium* | *simum* | 1 | 1 | 1 | 1 | 1 | 1 | 1 | 1 | 1 | 1 | 5 | 1800 | 9000 | 93 |
| Mammalia | *Lemur* | *catta* | 1 | 0 | 1 | 0 | 0 | 0 | 1 | 0 | 0 | 0 | 11 | 2.2 | 24.2 | 103 |
| Mammalia | *Cavia* | *porcellus* | 1 | 0 | 1 | 0 | 0 | 0 | 1 | 0 | 0 | 0 | 5 | 0.9 | 4.5 | 190 |

**Supplementary Table 7.** DNA extraction of 40 air samples collected in the Copenhagen Zoo in September and December. Samples collected with the 24 V sampler (MF) and the 5 V (SF) filters were set in PBS buffer (PBS). Further, the filters from MF were divided into 2 (A and B) during digestion. When possible, the two or three digests from the same sample were purified in the same column. This resulted in a total of 49 DNA extracts and 3 extraction negative controls.

| Sample number | Digests | Sample |
|---|---|---|
| 2 | NA | WV_Stable_30min_1 |
| 3 | NA | WV_Stable_30min_2 |
| 4 | NA | WV_Stable_Sampling blank |
| 5 | NA | WV_Stable_60min_1 |
| 6 | NA | WV_Stable_60min_2 |
| 7 | NA | WV_RainfHouse_Sampling blank |
| 8 | NA | WV_RainfHouse_30min_1 |
| 9 | NA | WV_RainfHouse_60min_1 |
| 10 | NA | WV_RainfHouse_30min_2 |
| 11 | NA | WV_RainfHouse_60min_2 |
| 12 | NA | WV_Outdoors_Sampling blank |
| 13 | NA | WV_Outdoors_30min_1 |
| 14 | NA | WV_Outdoors_30min_2 |
| 15 | NA | WV_Outdoors_60min_1 |
| 16 | NA | WV_Outdoors_60min_2 |
| 17 | NA | WV_Outdoors_30min_1 - September |
| 18 | NA | WV_Outdoors_30min_2- September |
| 19 | NA | WV_Outdoors_60min_1- September |
| 20 | NA | WV_Outdoors_60min_2- September |
| 21 | NA | Sterivex filter Blank |
| 22 | 22A | SF_Outdoors_30hrs_1 |
| | 22B | SF_Outdoors_30hrs_2 |
| 23 | 23A | SF_Stable_30hrs_2 |
| | 23B | PBS-SF_Stable_30hrs_2 |
| 24.1 | 24A | MF_Stable_5hrs_1 |

| | | |
|---|---|---|
| 24.2 | 24B | MS_OI_5_1 |
| | PBS | PBS-MS_OI_5_1 |
| 25.1 | 25A | MF_Stable_5hrs_2 |
| 25.2 | 25B | MF_Stable_5hrs_2 |
| | PBS | PBS-MF_Stable_5hrs_2 |
| 26.1 | 26A | MF_Stable_30min_1 |
| 26.2 | 26B | MF_Stable_30min_1 |
| | PBS | PBS-MF_Stable_30min_1 |
| 27 | 27A | MF_Stable_30min_2 |
| | 27B | MF_Stable_30min_2 |
| | PBS | PBS-MF_Stable_30min_2 |
| 28 | 28A | MF_Stable_60min_1 |
| | 28B | MF_Stable_60min_1 |
| | PBS | PBS-MF_Stable_60min_1 |
| 29 | 29.1 | MF_Stable_60min_2 |
| | 29.2 | MF_Stable_60min_2 |
| | 29B | PBS-MF_Stable_60min_2 |
| 30 | 30 | SF_Outdoors_30hrs_1 |
| | 30B | PBS-SF_Outdoors_30hrs_1 |
| 31 | 31 | SF_Outdoors_30hrs_2 |
| | 31B | PBS-SF_Outdoors_30hrs_2 |
| 32 | 32.1 | MF_Outdoors_60min_1 |
| | 32.2 | MF_Outdoors_60min_1 |
| | 32B | PBS-MF_Outdoors_60min_1 |
| 33 | 33.1 | MF_Outdoors_60min_2 |
| | 33.2 | MF_Outdoors_60min_2 |
| | 33B | PBS-MF_Outdoors_60min_2 |
| 34 | 34.1 | MF_Outdoors_30min_1 |
| | 34.2 | MF_Outdoors_30min_1 |
| | 34 B | PBS-MF_Outdoors_30min_1 |
| 35 | 35.1 | MF_Outdoors_30min_2 |
| | 35.2 | MF_Outdoors_30min_2 |

| | | |
|---|---|---|
| | 35 B | PBS buffer_MS_OO_30_2 |
| 36 | 36.1 | MF_Outside_5hrs_1 |
| | 36.2 | MF_Outside_5hrs_1 |
| | 36 B | PBS-MF_Outside_5hrs_1 |
| 37 | 37.1 | MF_Outside_5hrs_2 |
| | 37.2 | MF_Outside_5hrs_2 |
| | 37 B | PBS-MF_Outside_5hrs_2 |
| 38 | 38 | SF_RainfHouse_30hrs_1 |
| | 38 B | PBS-SF_RainfHouse_30hrs_1 |
| 39 | 39 | SF_RainfHouse_30hrs_2 |
| | 39 B | PBS-SF_RainfHouse_30hrs_2 |
| 40 | 40.1 | MF_RainfHouse_5hrs_1 |
| | 40.2 | MF_RainfHouse_5hrs_1 |
| | 40 B | PBS-MF_RainfHouse_5hrs_1 |
| 41 | 41.1 | MF_RainfHouse_5hrs_2 |
| | 41.2 | MF_RainfHouse_5hrs_2 |
| | 41 B | PBS-MF_RainfHouse_5hrs_2 |
| 42 | 42.1 | MF_RainfHouse_30min_1 |
| | 42.2 | MF_RainfHouse_30min_1 |
| | 42 B | PBS-MF_RainfHouse_30min_1 |
| 43 | 43.1 | MF_RainfHouse_30min_2 |
| | 43.2 | MF_RainfHouse_30min_2 |
| | 43 B | PBS-MF_RainfHouse_30min_2 |
| 44 | 44.1 | MF_RainfHouse_60min_1 |
| | 44.2 | MF_RainfHouse_60min_1 |
| | 44 B | PBS-MF_RainfHouse_60min_1 |
| 45 | 45.1 | MF_RainfHouse_60min_2 |
| | 45.2 | MF_RainfHouse_60min_2 |
| | 45 B | PBS-MF_RainfHouse_60min_2 |
| 46 | 46 | SF_Blank |
| | 46 B | PBS-SF_Blank |
| 47 | 47.1 | MF_Blank |

| | | |
|---|---|---|
| | 47.2 | MF_Blank |
| | 47 B | PBS-MF_Blank |
| 48 | 48 | PBS Blank |
| 49 | 49 | Room blank |
| Ex.bl1 | Ex.bl1 | Negative extraction control |
| Ex.bl2 | Ex.bl2 | Negative extraction control |
| Ex.bl3 | Ex.bl3 | Negative extraction control |

**Supplementary Methods:**

3D-blue prints of the housings used in the 24 V and 5 V particle samplers.

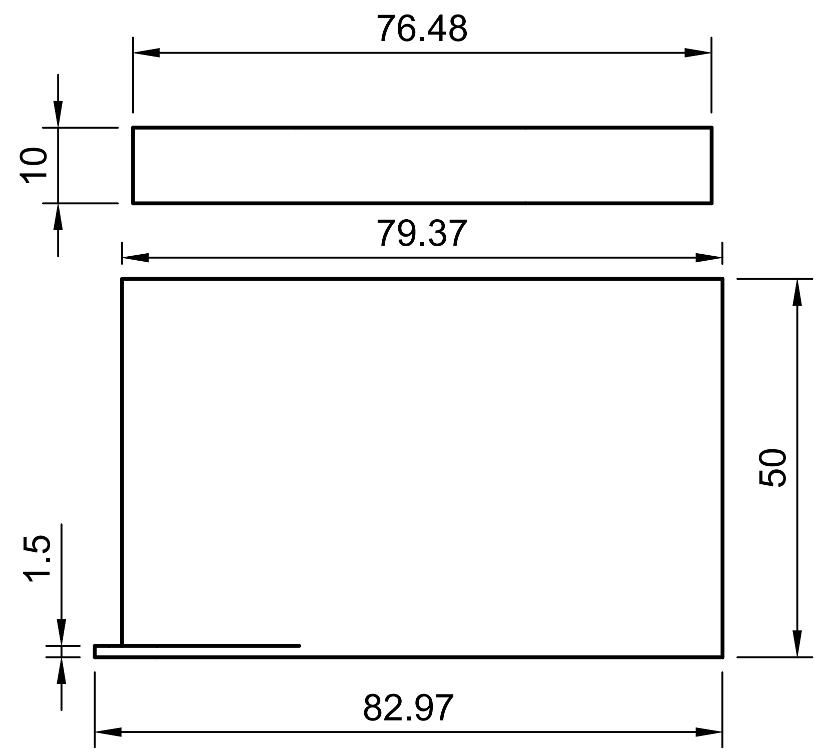


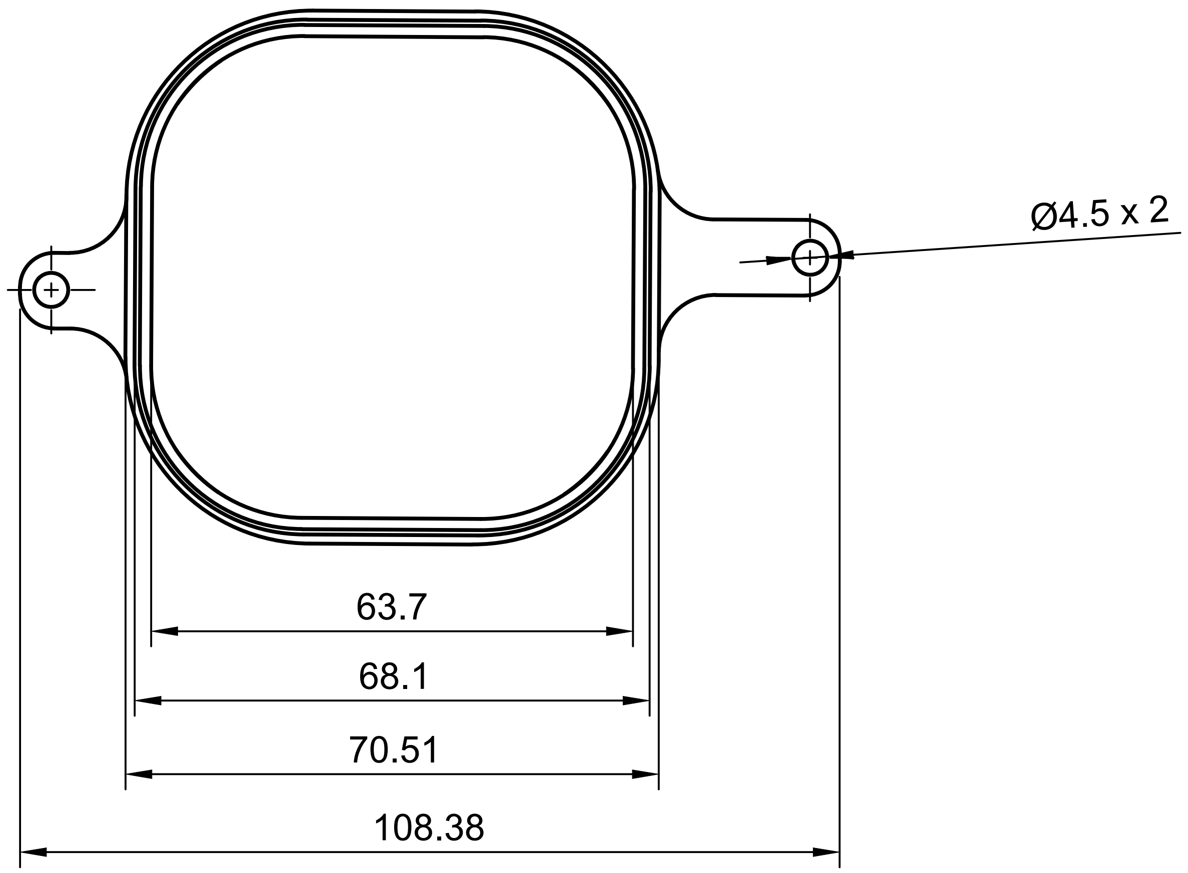


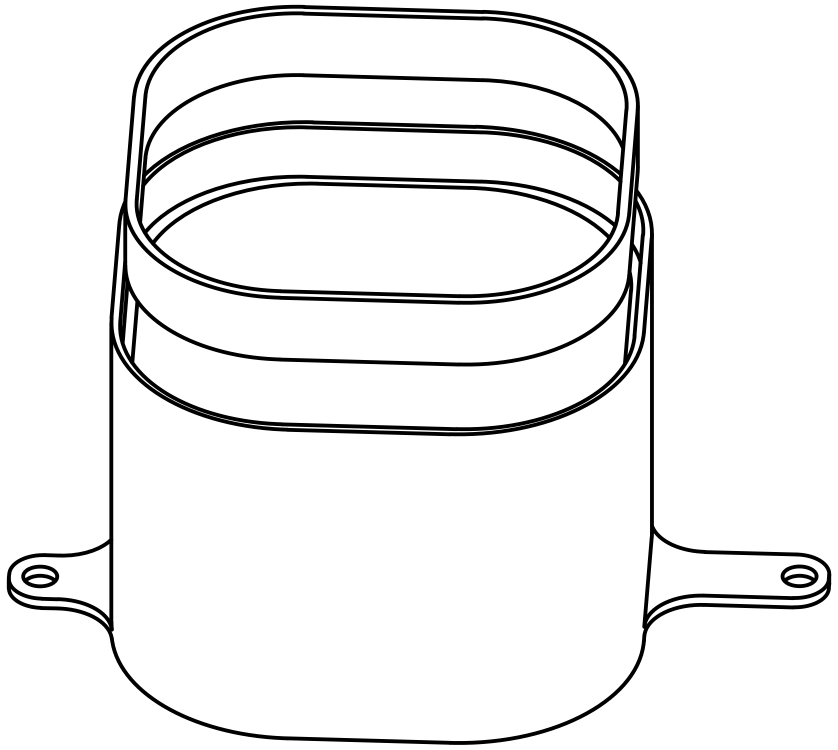

| Dept. | Technical reference | Created by AirLabs 11/05/2021 | Approved by | | |
|---|---|---|---|---|---|
| scale 1:1 | | Document type | Document status | | |
| | | Title Housing PM/fiber colector | DWG No. | | |
| | | | Rev. | Date of issue | Sheet 2/2 |

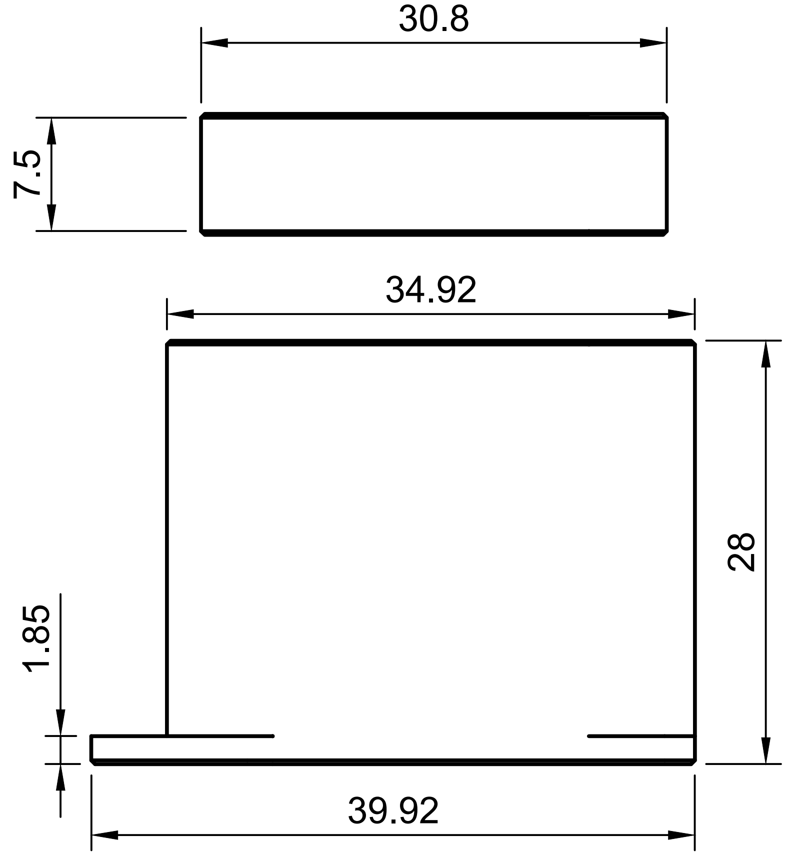


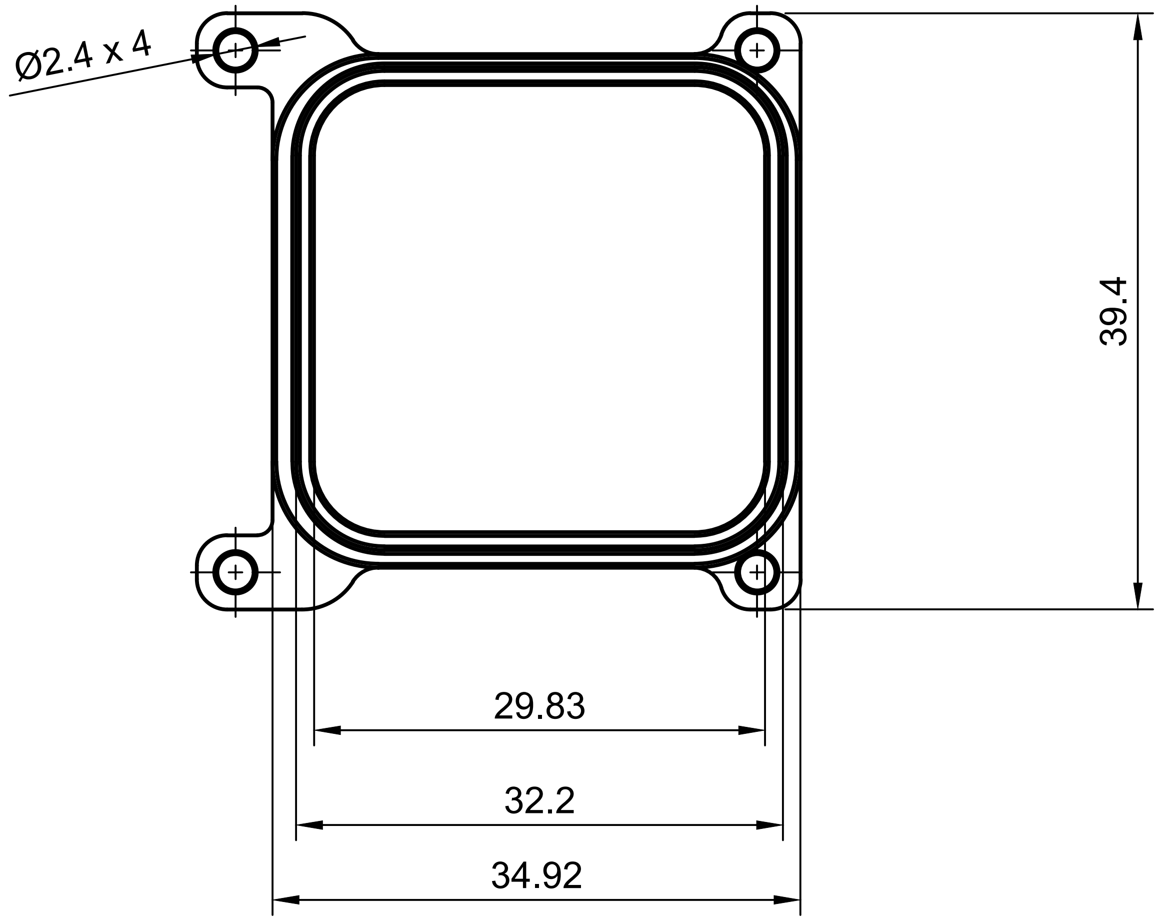


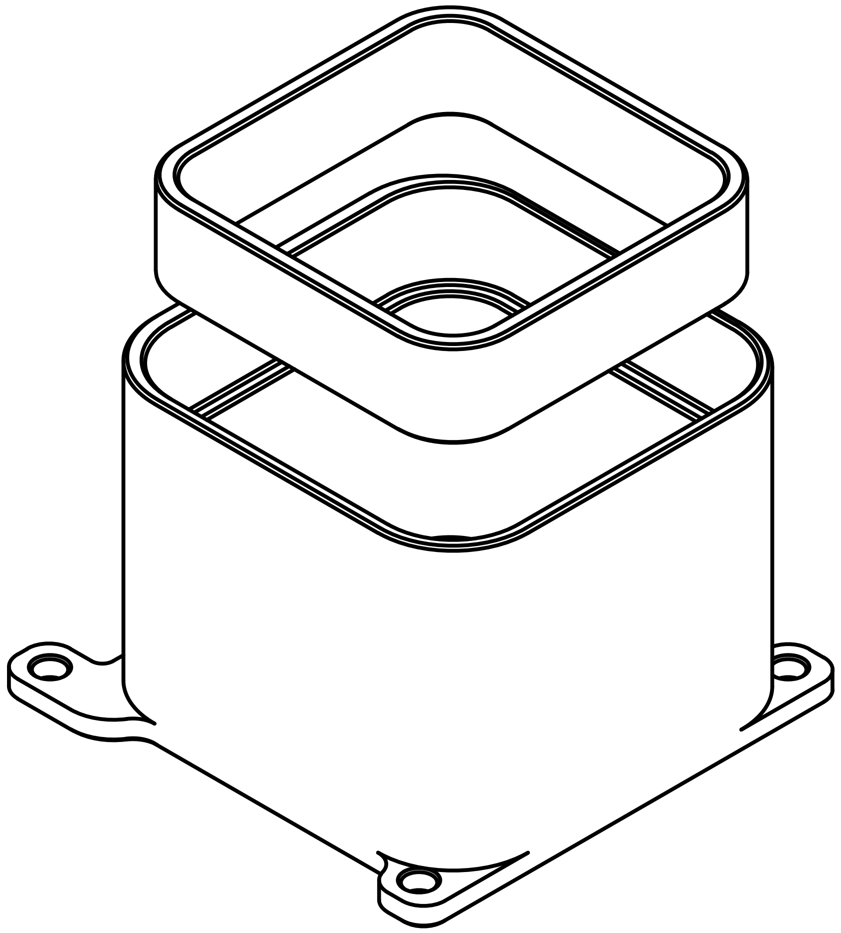

| Dept. | Technical reference | Created by AirLabs 11/05/2021 | Approved by |
|---|---|---|---|
| scale 2:1 | | Document type | Document status |
| | | Title **Housing** | DWG No. |
| | | | Rev. 1.0 / Date of issue / Sheet 1/1 |
